# Analysis of the worldwide diversity of *Xanthomonas hortorum* pv. *carotae*, the agent of bacterial blight of carrot, reveals two distinct populations

**DOI:** 10.1101/2023.04.21.537781

**Authors:** Enora Dupas, Karine Durand, Marie-Agnès Jacques

## Abstract

*Xanthomonas hortorum* pv. *carotae* is a pathogen responsible for the bacterial blight of carrot. While it infects carrot fields worldwide, its diversity remains unknown. Here, we validated by PCR the identification as *X. hortorum* pv. *carotae* of most of the strains isolated from carrot symptoms. We study their diversity by sequencing seven housekeeping genes. The analysis confirmed the identity of most of the strains previously identified as *X. hortorum* pv. *carotae* and highlighted the presence of two clusters inside the pathovar *carotae*. The 18 non-*X. hortorum* pv. *carotae* have been identified as *Xanthomonas* from other species. As the *X. hortorum* pv. *carotae* strains clustered only in two groups by this analysis, a new MLVA scheme was developed. It allowed to validate the subclusteration in two groups and observe diversity among them. At the same times based on this study and data from the literature, we proposed the strain CFBP 7900 as the neopathotype strain of the pathovar. The strain should replacement the holopathotype strain CFBP 4997, as it is no longer available in international collection.

## 2. Introduction

Global food security is threatened by the dispersal of plant diseases that are favored by the increase of plant material trade and transport, crop intensification and climate change (Savary and Willocquet, 2020). Several recent disease emergence or reemergence highlight the severe impact such unanticipated events may have on socio-economic development of large areas (Chapman et al., 2017; Pratt et al., 2017; Schneider et al., 2020). Better prevention and control of plant pathogens is essential to ensure sufficient plant production and depends on a good understanding of pathogen diversity and population genomic surveillance (Ristaino et al., 2021). The use of efficient molecular tools to type pathogen populations is nowadays a critical step to develop adapted management and control strategies for plant diseases as they may inform resistance breeding and reveal the source of the outbreak strains.

*Xanthomonas hortorum* pv. *carotae* is a seed-borne pathogen (Umesh et al., 1998) responsible for bacterial blight of carrot (Kendrick, 1934). The first description of symptoms of carrot leaf blight dates back to 1855 when Kühn et al. described it in Germany (Farrar et al., 2004). Then the bacterium was detected in 1931 in California by Kendrick who named it *Pseudomonas carotae* (Kendrick, 1934). Then, in 1939 Dowson proposed a novel taxonomic framework and reclassified all *Pseudomonas* isolated on plants that form yellow colonies in a new genus named *Xanthomonas* (Dowson, 1939). But it was not until 1978, following the creation of the term “pathovar” in bacterial taxonomy, that it was included in the *Xanthomonas* campestris species as the pv. *carotae* (Young et al., 1978), and finally transferred to the species X. *hortorum* based on DNA/DNA hybridizations (Palleroni et al., 1993). Bacterial blight is a common disease of carrot wherever carrot is cropped. The most recent reports concerned Mauritius, Indiana state in the USA, Korea and Spain (Pruvost et al., 2010; du Toit et al., 2014; Myung et al., 2014; Palomo et al., 2021).

On carrot plants infected by *X. hortorum* pv. *carotae*, the disease develops as small, brown or dark, irregular, greasy, water-soaked spots surrounded by a yellow chlorotic halo that evolve in necrosis (du Toit et al., 2005; Kimbrel et al., 2011). Leaflets may curl up and become brittle as if they had been burned. The whole carrot, from the root to the umbel, can be infected by the bacterium and affected by the disease, preventing them from being mechanically harvested by gripping the leaves (Farrar et al., 2004) and causing significant yield losses (Christianson et al., 2015). Moreover, all aerial organs of carrot are susceptible to leaf blight, and can be infected by the bacterium leading to seed-infection, even in the absence of symptoms (du Toit et al., 2005). The incidence of the disease is linked to the initial inoculum size on seeds and is highly dependent on climatic conditions (Umesh et al., 1998). Besides being seed-transmitted, this pathogen is disseminated by aerosols formed during rain, irrigation or field operations, and infected crop residues (du Toit et al., 2005; Christianson et al., 2015).

The complete genome of a *X. hortorum* pv. *carotae* strain (CFBP 7900 = M081), isolated in 2008 from infected carrot seeds, was sequenced by Kimbrel and colleagues (2011) (Kimbrel et al., 2011). This 5.1 Mbp genome sequence is an invaluable resource for studying the pathovar *carotae*, searching for specific signatures using comparative genomics to gain insights in pathogenicity and develop diagnostics tools. The genomic sequence of the strain M081 showed that *X. hortorum* pv. *carotae*, like most Gram-negative plant pathogens, presents all six secretion systems that allow the bacterium to interact with its biotic and abiotic environment (Kimbrel et al., 2011). In particular, this plant pathogen relies on a classical *Xanthomonas* type three secretion system (T3SS) to deliver a range of T3 effectors in host cell, some of which were demonstrated to elicit effector-triggered immunity (Kimbrel et al., 2011).

Being a seed-pathogen, efforts have been devoted to develop seed-health tests. The most recent International Seed Testing Association-validated protocol (ISTA, 2023) recommend a seed wash dilution-plating assay followed by a PCR test (Meng et al., 2004) to confirm the identity of suspect isolates. The International Seed Federation protocol (ISHI-Veg, 2021) proposes to replace the gel-based Meng’s PCR test in the ISTA protocol by a TaqMan test combining primers and probes from Barnhoorn 2014 (MVS*Xhc*3 set) and from Temple *et al*. 2013 (*Xhc*-q2 set) (Temple et al., 2013; Barnhoorn, 2014).

While MultiLocus Sequence Analysis (MLSA) is mostly used to infer phylogenetic relationships among species within a genus and species delineation, i.e. for taxonomic purposes (Gevers et al., 2005; Glaeser and Kämpfer, 2015), MultiLocus Sequence Typing (MLST) is devoted to the characterization of isolates (Maiden et al., 1998). Both methods rely on the sequencing of internal fragments of 4 to 7 housekeeping genes. Sequences are generally concatenated and used for comparative analyses in MLSA, while in MLST sequences obtained for a given taxon are compared and different sequences are assigned as distinct alleles and, for each isolate, the alleles at each of the seven loci define the allelic profile or sequence type (ST) that is used for subsequent analyses. These two methods have proven highly relevant for analyses at the species or sub-species level, but generally fail to provide sufficient resolution for infra-subspecific levels and in particular for highly clonal organisms (Maiden, 2006; Ferreira et al., 2019; Dupas et al., 2023). Multilocus Variable Number of Tandem Repeats (VNTR) analysis (MLVA) is a high-resolution method for monitoring epidemic clones and assessing population structure for infra-subspecific bacterial taxa. A typical VNTR locus shows large range of copy numbers even among highly related bacterial strains (Jackson, 2010). MLVA has been used as a powerful tool for outbreak detection and source tracing of various important plant pathogenic bacteria, such as *Xylella fastidiosa* (Dupas et al., 2023), *Xanthomonas citri* pv. *citri* (Leduc et al., 2015); *Erwinia amylovora* (Bühlmann et al., 2014) and *Pseudomonas syringae* pv. *actinidiae* (Cunty et al., 2015). In plant pathology, a specific taxon of invaluable importance is the pathovar. A pathovar groups strains that are pathogenic on the same host range, on which they induced the same symptomatology (Dye et al., 1980). The name-bearer of a pathovar is a pathotype strain, by similarity with the type strain of a species (Dye et al., 1980). This pathotype strain presents all biological characteristics of the taxon that has been described. *X. hortorum* pv. *carotae* was described by Kendrick et al., in 1934 to group oxidative strains forming yellow colonies on nutrient agar and that are responsible for bacterial blight of carrot. In contrast, the currently stored specimens of the pathotype strain of *X. hortorum* pv. *carotae* (CFBP 4997 = ICMP 5723 = LMG 8646 = NCPPB 1422) are copies of a fermentative strain belonging to the *Enterobacter* genus. This indicates that a contaminant was distributed and stored instead of the original pathotype strain, and thus justifying the designation of a novel pathotype strain following rule 9.4 of the Committee on the Taxonomy of Plant Pathogenic Bacteria of the ISPP (Dye et al., 1980).

The objective of this study was to assess the genetic diversity of a collection of *X. hortorum* pv. *carotae* strains isolated worldwide over two decades. First, we identified the strains using a PCR test and phylogenetically assigned the strains based on concatenated sequences of seven housekeeping genes (MLSA). This part of the work allowed us to propose a neopathotype strain to X. *hortorum* pv. *carotae*. We then developed an 8-VNTR scheme to assess the genetic structure of this pathovar to inform genetic resistance studies of carrot to this pathogen and trace the links among strains.

## 3. Methods

### 3.1. Bacterial strains and growth conditions

A collection of 140 *Xanthomonas*-like strains isolated in Canada, France, Germany, the Netherlands, Mauritius island, and the USA between 1994 and 2012 from carrot fields or carrot seeds (*Daucus carota*), and weeds was used in this study (Table 1). Among them, seven strains were isolated from symptomless weeds sampled in *X. hortorum* pv. *carotae* -infected carrot fields in France. All strains were preserved by the French Collection of Plant-Associated Bacteria (CIRM-CFBP, INRAE, https://doi.org/10.15454/E8XX-4Z18). Strains were grown on TSA 100% medium (tryptone 1,7g/L, soy peptone 0,3 g/L, NaCl 0,25 g/L, K2HPO4 0,5 g/L, agar 0,5 g/L, Polyflor 2 mL/L, adjusted at pH 7,2) at 28°C for 48h. Bacterial suspensions were prepared from fresh culture in sterile distilled water and adjusted at 1×10^8^ CFU/mL. Bacterial suspensions were stored at −20°C prior analysis.

**Table 1:**
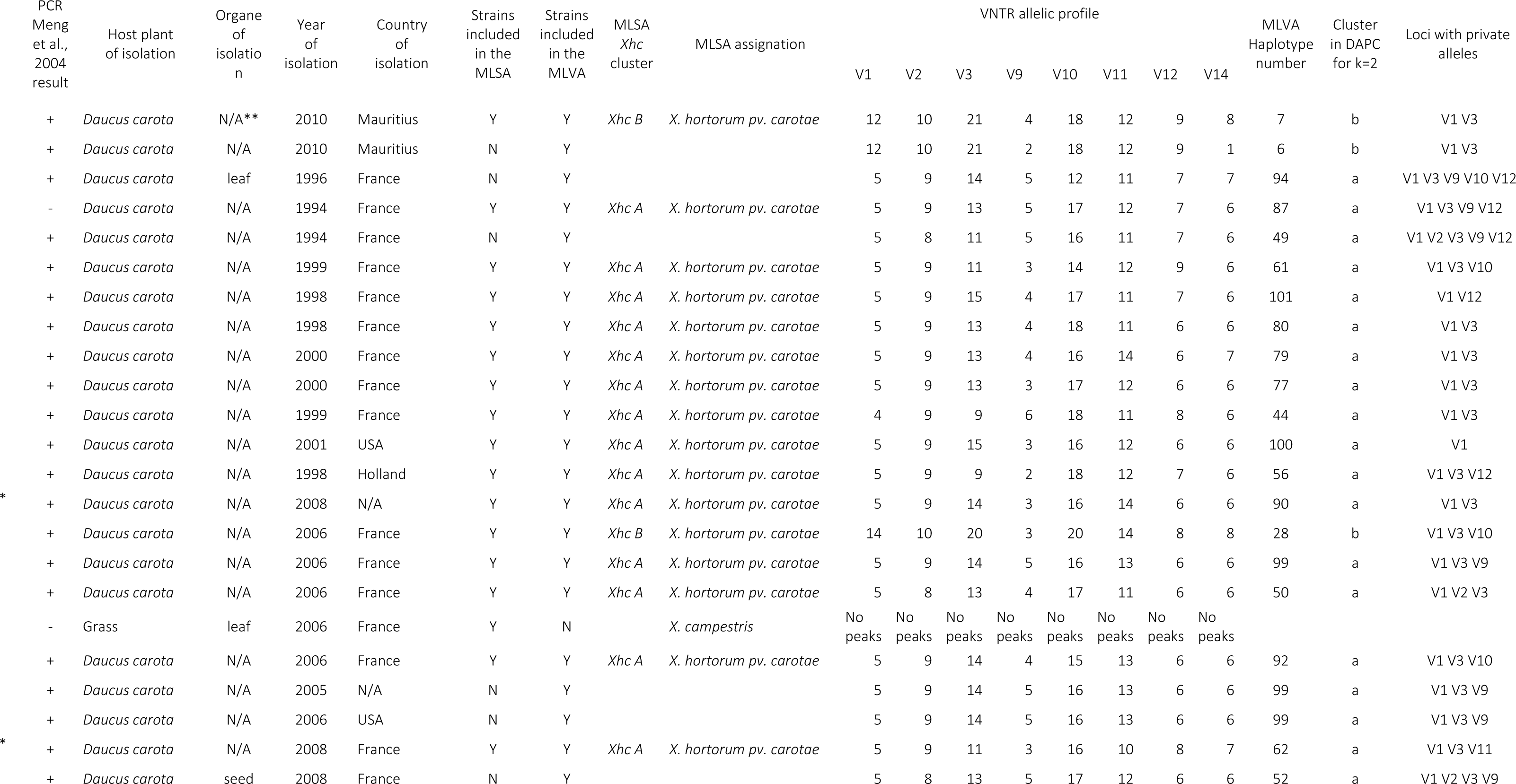
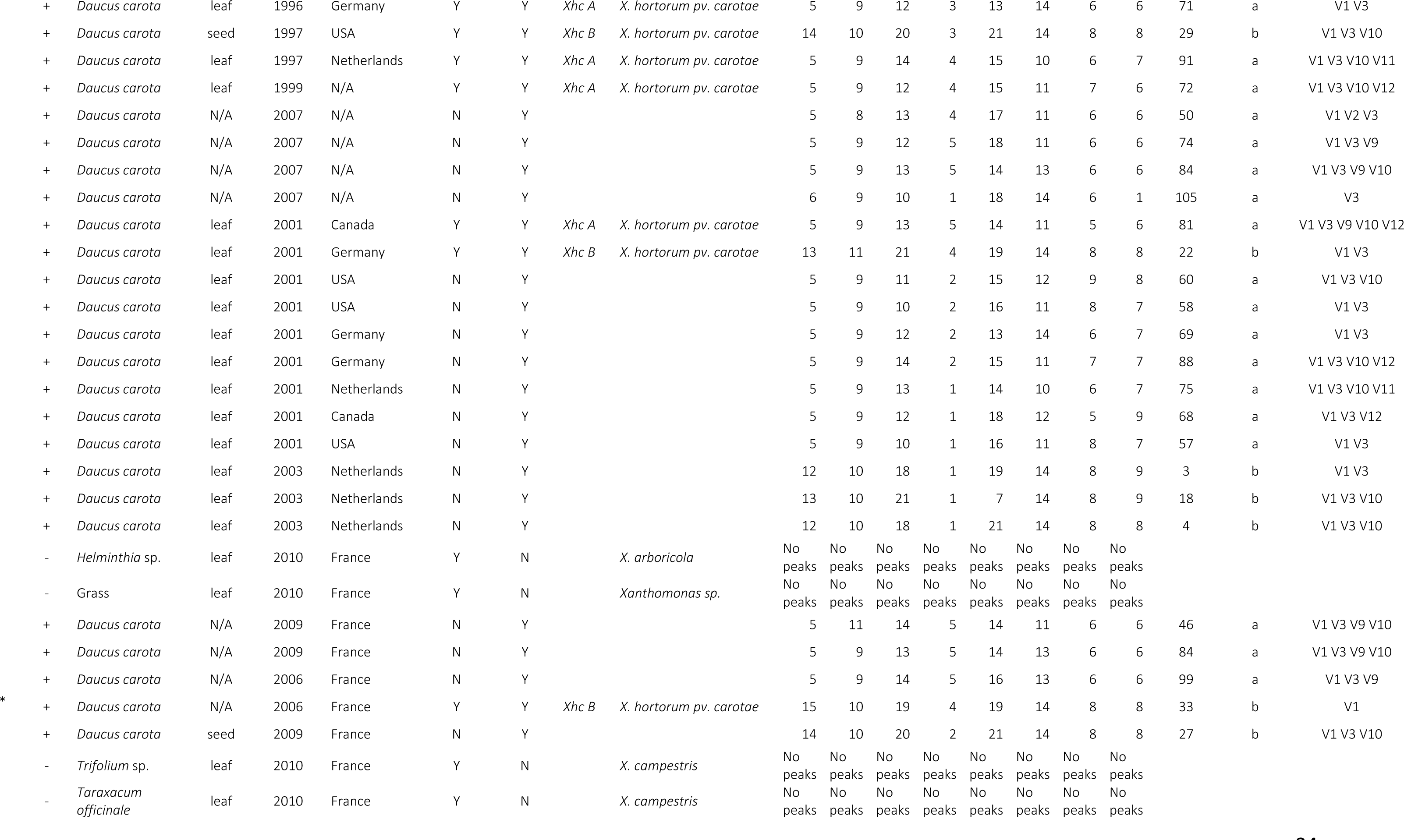
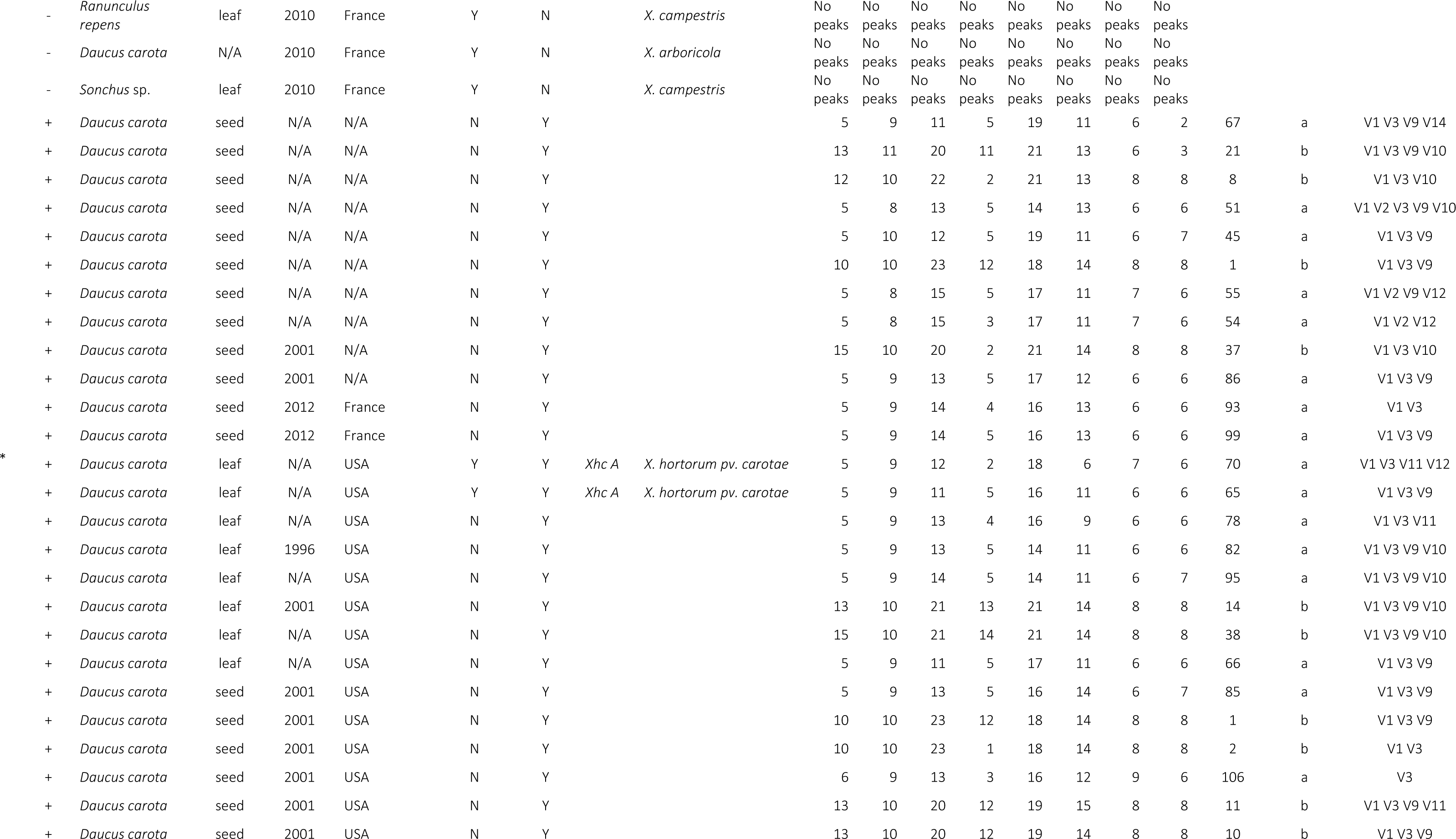
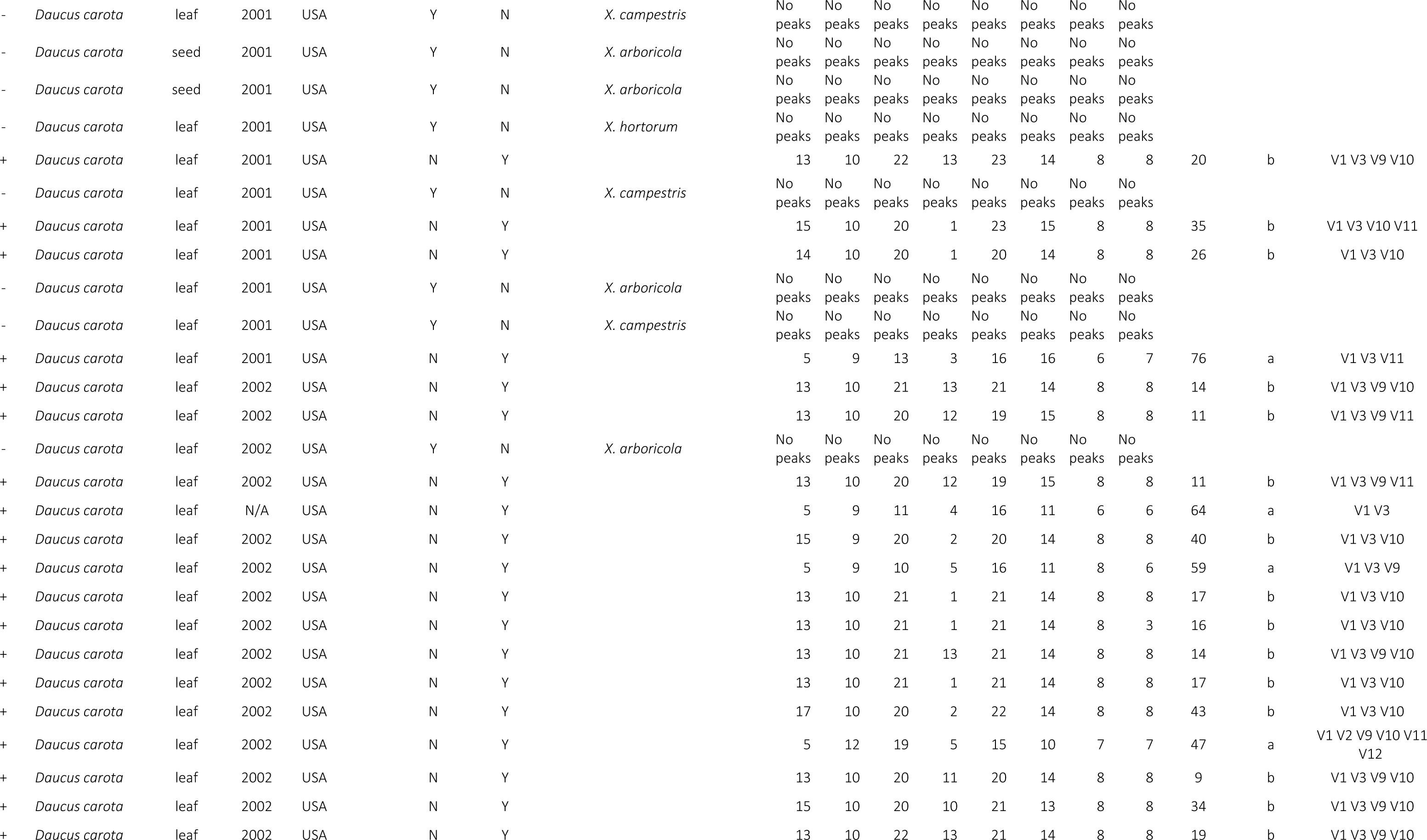
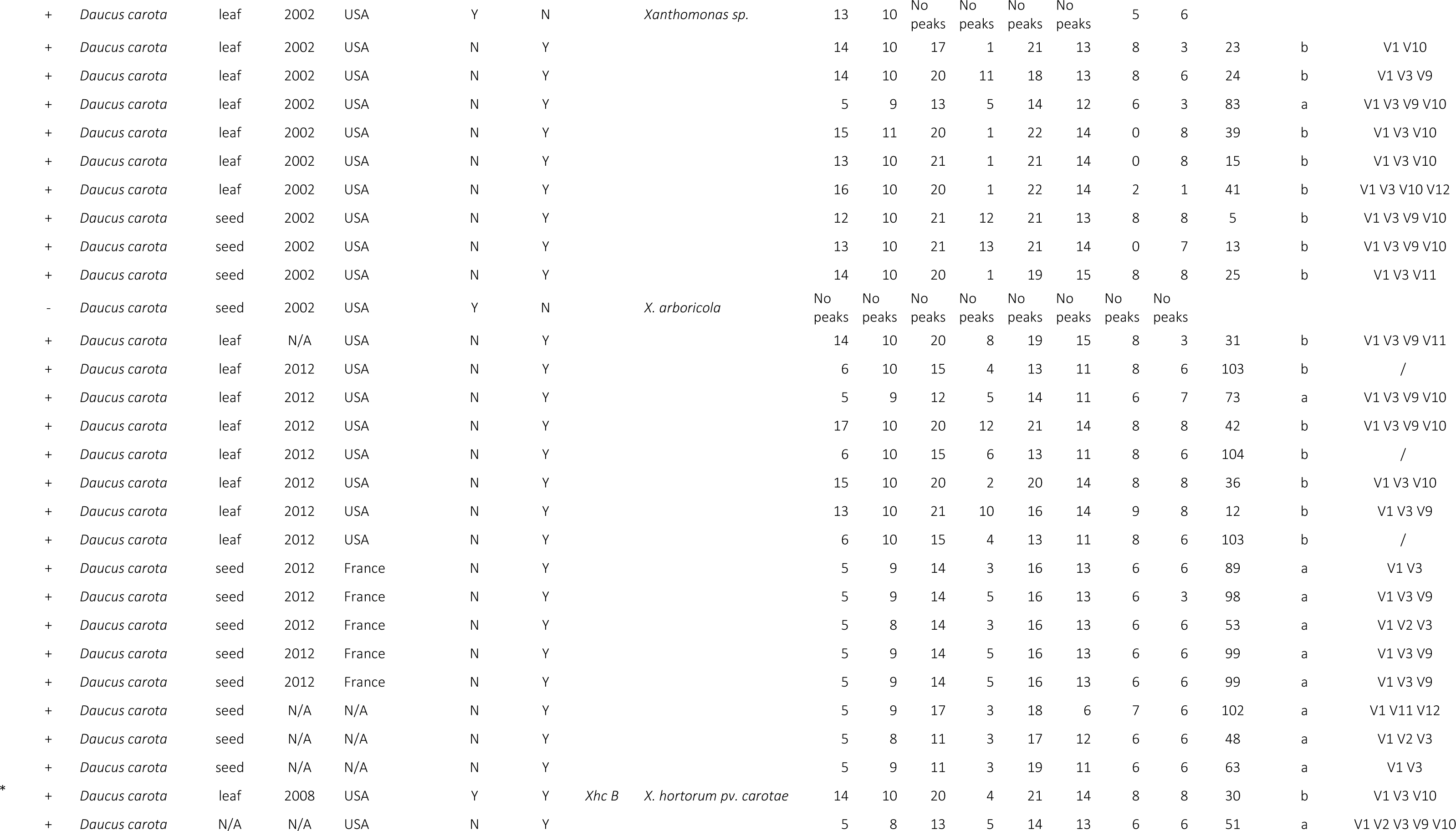
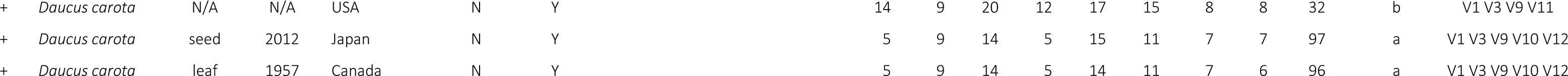
List of the 140 strains used in this study. * strains analyzed to select the VNTR list of the study, ** not available

### 3.2. Strain identification

*X. hortorum* pv. *carotae* -like identity was validated using the *X. hortorum* pv. *carotae* specific PCR test designed by Meng *et al*., (2004). PCR was performed following authors’ protocol on bacterial suspensions. PCR products were revealed in 1.5% agarose gel in 1X Tris-acetate-EDTA buffer. Gels were stained with ethidium bromide (2 mg/L) and amplified fragments visualized with ultraviolet light. The identity of the strains as *X. hortorum* pv. *carotae* was confirmed if an amplicon of 350 bp was visible on the gel.

### 3.3. Multi Locus Sequence Analysis (MLSA) of *X. hortorum* pv. *carotae*

The diversity of a subcollection of 44 strains (Table 1), chosen for their diversity of years and places of isolation, was analyzed using a classical MLSA scheme based on fragments of seven housekeeping genes (*atpD, dnaK, efP, fyuA, glnA, gyrB* et *rpoD*) previously designed for *Xanthomonas* spp. by Mhedbi-Hajri et al., (2013) (Table 1). PCRs were performed following authors’ protocol on bacterial suspensions. The PCR products were sent to GenoScreen (France) for both forward and reverse sequencing. For each strain, forward and reverse sequences were assembled and nucleotide assignation checked using Geneious 9.1.8 software (Biomatters). Consensus sequences were aligned and trimmed using BioEdit software (Hall, 1999). Maximum of likelihood trees were constructed using a bootstrap method of 100 to 1, 000 replications and Kimura 2-parameter model with the Mega 11 software (Tamura et al., 2021). Gene sequences from the type strains of 34 *Xanthomonas* species and seven pathotype strains of *X. hortorum* pathovars were added to the analyses (Table S1).

### 3.4. Design and development of the 8-VNTRs scheme

The complete genome sequence of the strain M081 (=CFBP 7900) of *X. hortorum* pv. *carotae* isolated from *Daucus carota* seeds in 2008 in Oregon (USA), was screened for the presence of tandem repeat (TR) loci using https://bioinfo-web.mpl.ird.fr/xantho/utils/ set with default parameters. A couple of primers was designed for the 14 identified VNTR loci on the flanking region of TR loci using Primer3 (Koressaar and Remm, 2007). The existence of polymorphism was checked by PCR on a subset of six *X. hortorum* pv. *carotae* strains (Table 1). PCRs were performed in a final volume of 20 µL containing 1X of Colorless GoTaq® Flexi Buffer (Promega), 0.0625 mM of each dNTP, 0.125 µM of each primer, 0.4U of GoTaq® Flexi DNA Polymerase (Promega), 1.5 mM of MgCl2 (Promega) and 5 µL of bacterial suspension. PCR amplifications were performed using the following cycling program: denaturation step of 5 min at 95°C, followed by 32 cycles of denaturation of 30 sec at 94°C, annealing of 30 sec at 60 or 55°C (depending of the VNTR, see Table 2, Table S2) and extension of 30 sec at 72°C, and a final extension step of 10 min at 72°C. PCR products were revealed and check for diversity in an agarose gel using the same protocol as for §3.2.

**Table 2:**
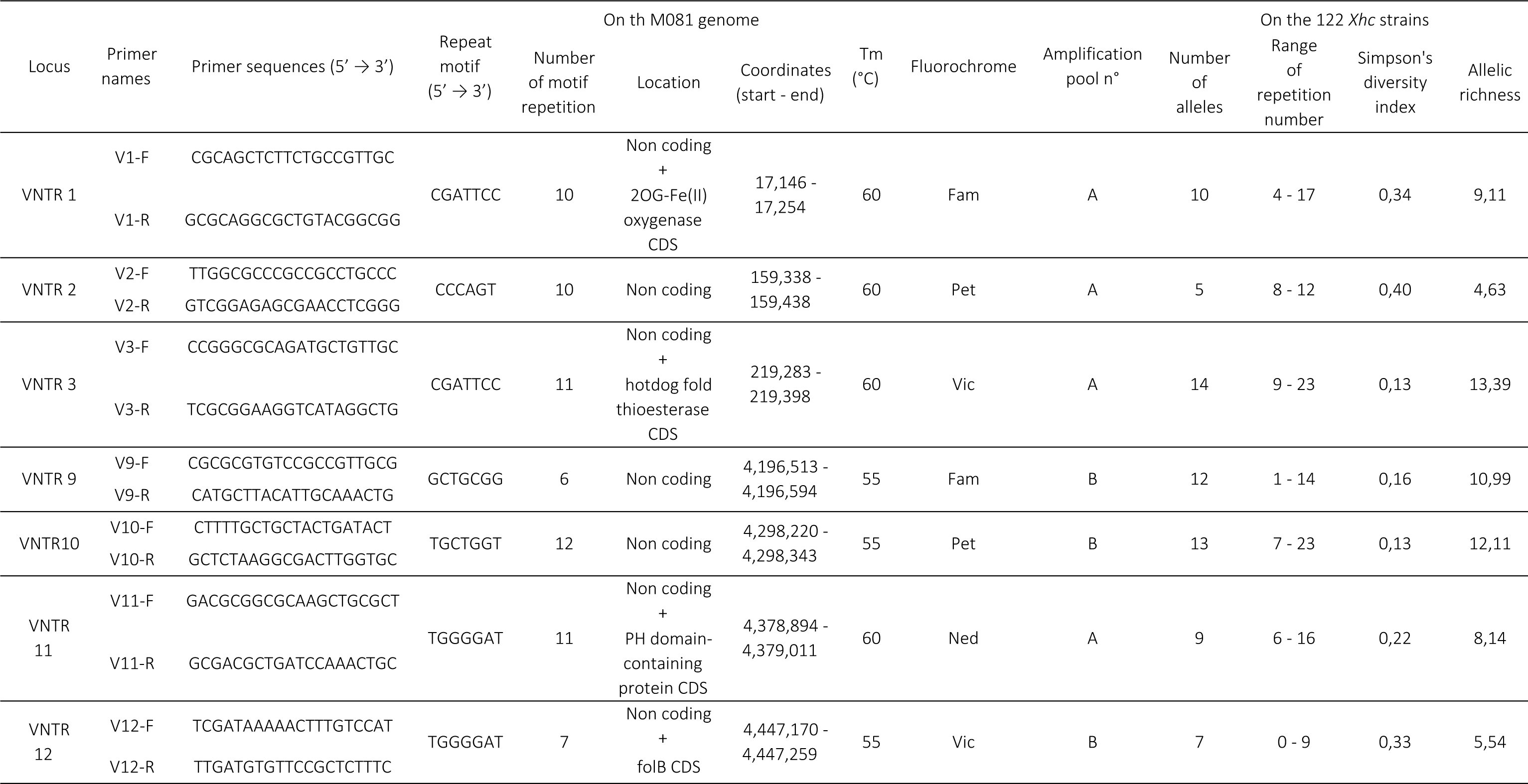
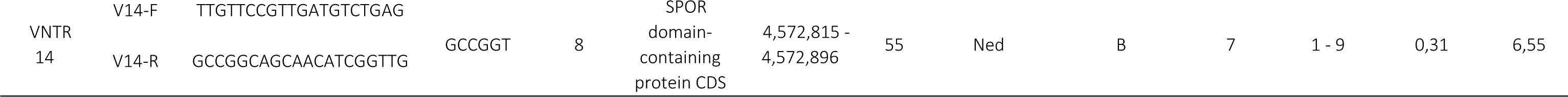
Nomenclature, location, function and genetic diversity of the 8 TR loci.

The eight forward primers showing diversity (V1, V2, V3, V9, V10, V11, V12 and V14) were labeled with a fluorescent dye (6-FAM, VIC, NED or PET, Table 2). The labeled primers were used to amplify VNTRs within the entire *X. hortorum* pv. *carotae* collection (Table 1) using the same reaction mix and amplification protocol as used to test the primers. Post-amplification, PCR products were multiplexed in order to obtain a solution composed of the four dyes. 1 µL of each PCR product labeled with 6-FAM, VIC, NED and NED dye, 2 µL of product labeled with PET dye and 17 µL of distilled water were mixed. Then 2.4 µL of multiplexed PCR products were mixed with 9.35 µL of Hi-Di formamide (Sigma-Aldrich) and 0.15 µL of Genescan 500 Liz internal line size standard (Applied Biosystems) for capillary electrophoresis analysis, using an ABI PRISM 3130 genetic analyzer (Applied Biosystems): injection of 16 sec at 1.2 kvolts, voltage steps 20 nk, interval of 15 sec, data delay time of 60 sec, run of 12000 sec at 15 kvolts.

### 3.5. Data scoring

Electrophoregrams were analyzed using Geneious 9.1.8 software (Biomatters). Ladder and VNTR peaks were automatically detected using the predict peaks mode and carefully checked by eye to correct artefacts or small miss-annotated peaks. The number of repeats in each locus was calculated based on the fragment sizes using Geneious 9.1.8 software (Biomatters).

### 3.6. MLVA analyses and statistics

Minimum spanning trees (MST) were drawn using BioNumerics 7.6 (Applied Maths). Clonal complexes grouped haplotypes differing by a maximum of two TRs. The discriminatory power of the MLVA was calculated using http://insilico.ehu.es/mini_tools/discriminatory_power/index.php. Assignment of samples to a cluster was performed by Discriminant Analysis of Principal Components (DAPC) on 20 independent k-means runs in order to verify the stability of the clustering. Analyses were performed using a single individual per haplotype with the adegenet package from the R software (Jombart, 2008). Simpson’s index of diversity using BioNumerics 7.6 (Applied Maths) and allelic richness using Fstat 2.9.4 (Goudet, 2005) were calculated to access the discriminatory power of each VNTR.

### 3.7. Looking for pathogenicity related genes

The presence of the three genes *hrcC, hrcV* and *hrcN* coding core proteins of the T3SS was search for in bacterial genomes using primers and protocol designed by Hajri et al. (2011) (Hajri et al., 2011). PCR products were revealed in 2% agarose gel in 1X Tris-acetate-EDTA buffer. Gels were stained with ethidium bromide and amplified fragments visualized with ultraviolet light. The putative presence of a functional T3SS was indicated by the co-occurrence of amplicons of 575 bp for *hrcC*, 524 bp for *hrcN* and 367 bp for *hrcV* on gels.

### 3.8. Hypersensitive response (HR) on tobacco

The 18 strains that did not amplified using the *X. hortorum* pv. *carotae* -specific PCR test (Meng et al., 2004) were inoculated on tobacco (*Nicotiana benthamiana*). Two *X. hortorum* pv. *carotae* strains (CFBP 7768 and CFBP 7900) were used as positive controls. Suspensions at 1×10^8^ CFU/mL were infiltrated into the leaf limbs of tobaccos with a needleless syringe. Negative control plants were infiltrated with sterile distilled water. The presence of symptoms was observed at 48h and confirmed at 72h post-inoculation.

### 3.9. Pathogenicity on carrot

Pathogenicity of the 18 strains that did not amplify using the *X. hortorum* pv. *carotae* -specific PCR test (Meng et al., 2004) was tested by inoculation on 6 weeks-old plants of *Daucus carota* cv. Nansen grown in greenhouse. Strains CFBP 7768 and CFBP 7900 were used as positive controls and water as negative control. Inoculations were performed by soaking carrot foliage at the 3 leaf-stage in a suspension at 1×10^7^ CFU/mL during 30 sec. Inoculated plants were placed in mini-tunnels and maintained under saturated hygrometry during 48h. The assay was performed in duplicate and symptoms were observed three weeks after inoculation.

## 4. Results

### 4.1. Preliminary identification of the strain

The 140 strains used in this study were all isolated from carrot leaves, carrot seeds, or from weeds sampled in *X. hortorum* pv. *carotae* -infected carrot fields. All strains presented a colony morphology characteristic of *Xanthomonas* on TSA 10% medium, being yellow and butyrous. We used the *X. hortorum* pv. *carotae* Meng’s PCR test (Meng et al., 2004)) to identify these strains, as this test is known to be highly specific (ISTA, 2023). A correct signal was obtained for 122 strains (Table 1). These 122 strains were all isolated from carrot seeds or plants, while the 18 remaining ones were isolated from carrots (11 strains) or weeds (7 strains), meaning that no strains from weeds were identified as *X. hortorum* pv. *carotae* (no amplification in at least three different PCR runs).

### 4.2. Taxonomic assignation of the strains

To confirm preliminary identification provided by the use of the Meng’s *X. hortorum* pv. *carotae* -specific PCR test, a subset of 44 strains maximizing the diversity in terms of host and places of isolation (Table 1) was phylogenetically assigned by MLSA. This subset included 26 strains that tested positive and the 18 strains testing negative with the *X. hortorum* pv. *Carotae* -specific Meng’s test. The strain CFBP7900 (=M081) was included in this subset as it is known to be pathogenic on carrot. The seven housekeeping genes were amplified in all selected 44 strains. Twenty-five of the 26 strains preliminary identified as *X. hortorum* pv. *carotae* grouped in two clusters (Figure 1). These two clusters differed on 65 positions out of the 4,189 bp analyzed and are supported by an 79% bootstrap value. The cluster A grouped 20 strains including the strain CFBP 7706 that did not give any signal with the *X. hortorum* pv. *carotae* Meng’s test. This cluster was supported by a 100% bootstrap value and grouped 18 strains having 100% identical MLSA sequences while the 2 others (CFBP 7760 and CFBP 7762) differed by only one nucleotide on the *gyrB* gene fragment (A <-> C bp n°3612/4189 of the final concatenate dataset). The cluster B grouped 6 strains and the sequences retrieved from the genome sequence of the strain M081, with a 100% identical sequence for the 7 housekeeping gene fragments and a bootstrap value of 100.

**Figure 1:**
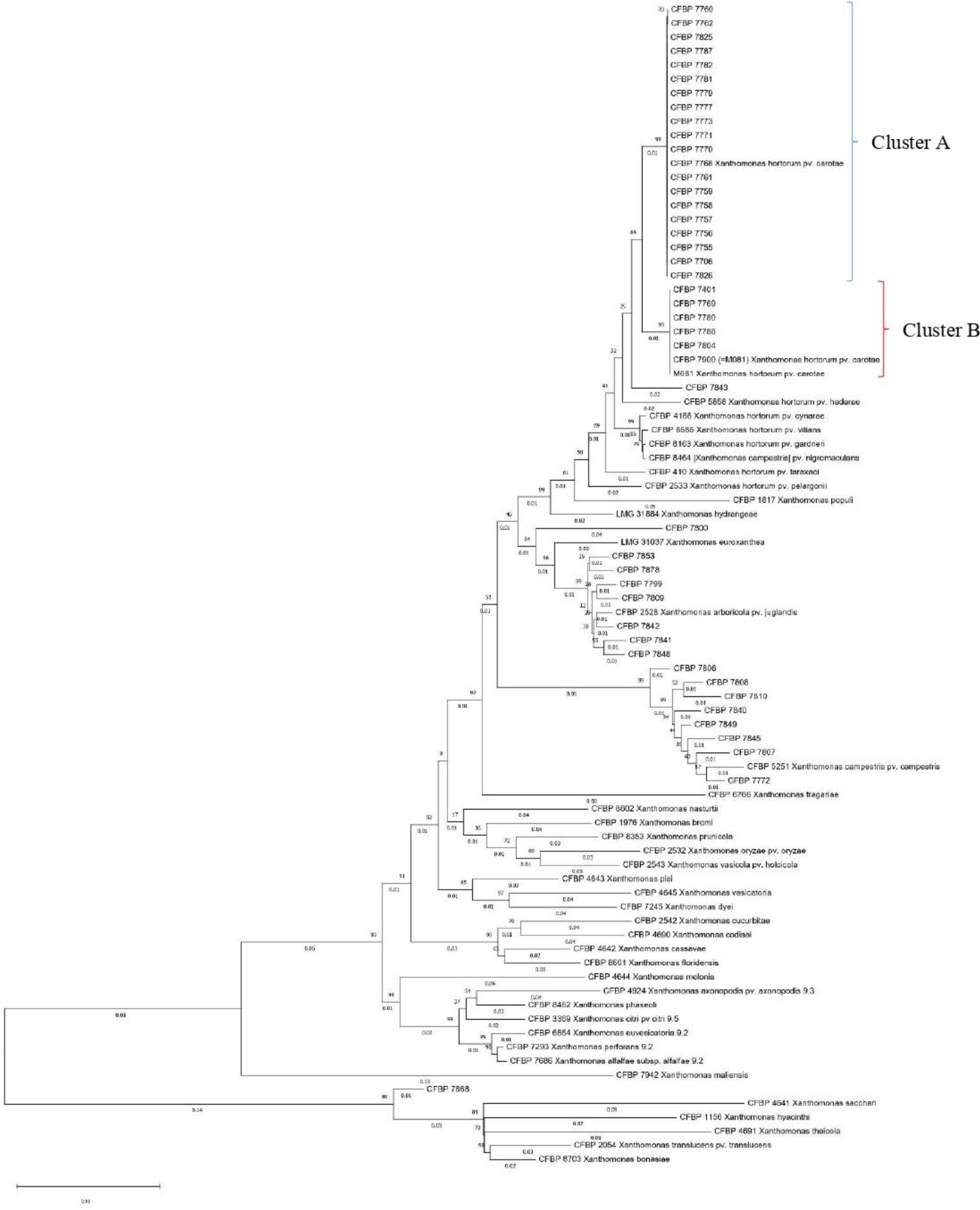
Phylogenetic relationships of the 26 *X. hortorum* **pv.** *carotae* and 18 non-*X. hortorum* pv. *carotae* strains from our collection evaluated by MLSA and compared to the pathotype strains of *X. hortorum* and and the type strains of all species currently included in the genus *Xanthomonas*. Maximum likelihood tree was constructed on concatenated portions of *atpD, dnaK, efP, fyuA, glnA, gyrB* and *rpoD* genes (for a total length of 4,205 bp) with bootstrap scores calculated on 100 replicates.

These 26 *X. hortorum* pv. *carotae* strains were genetically distant from the other 18 strains, 17 of which tested negative with the *X. hortorum* pv. *carotae* Meng’s test and one (strain CFBP 7868) giving an amplification using the *X. hortorum* pv. *carotae* Meng’s test (Figure 1). These strains were dispersed within the *Xanthomonas* genus. The strain CFBP 7843 grouped within the species X. *hortorum*, while the others are more distantly related (Figure 1). The strains CFBP 7799, CFBP 7809, CFBP 7841, CFBP 7842, CFBP 7848, CFBP 7853 and CFBP 7878 grouped within the *X. arboricola* species, the strains CFBP 7772, CFBP 7806, CFBP 7807, CFBP 7808, CFBP 7810, CFBP 7840, CFBP 7845 and CFBP 7849 grouped within the *X. campestris* species. The strains CFBP 7868 and CFBP 7800 could not be assigned to any species, being at the root of the genus or to loosely related to any other species. Interestingly, the strain CFBP 7868 grouped within the *X. hortorum* pv. *carotae* pathovar looking only at the *dnaK* ML tree, but was at the root of the six other housekeeping gene trees (Fig S1 to S7).

### 4.3. Pathogenicity of the strains

Pathogenicity of the 17 strains that tested negative with the Meng’s test plus strain CFBP 7868, which did not cluster within *X. hortorum* pv. *carotae,* was tested on *D. carota* cv. Nansen by soaking the three first leaves of 6 weeks-old plants in 1 × 10^7^ cfu/ml bacterial suspensions. Strains CFBP 7768 and CFBP 7900 were used as positive controls and represented clusters A and B, respectively. We also tested the strain CFBP 7706, that did not amplify using Meng’s test but grouped in the *X. hortorum* pv. *carotae* cluster A by MLSA (Figure 1). This strain induced typical symptoms of carrot bacterial blight, similar to those of the positive controls, 17 days post-inoculation. Strains CFBP 7800, CFBP 7842, CFBP 7843, CFBP 7853 did not cause any symptoms on carrot. Small white dry spots, typical of a hypersensitive reaction (HR), were clearly visible on carrot leaves inoculated with the 14 other strains (CFBP 7772, CFBP 7799, CFBP 7806, CFBP 7807, CFBP 7808, CFBP 7809, CFBP 7810, CFBP 7840, CFBP 7841, CFBP 7845, CFBP 7848, CFBP 7849, CFBP 7868, CFBP 7878), whatever their hosts of isolation, *i.e.* carrot or weeds (Table 3, Figure 2).

**Figure 2:**
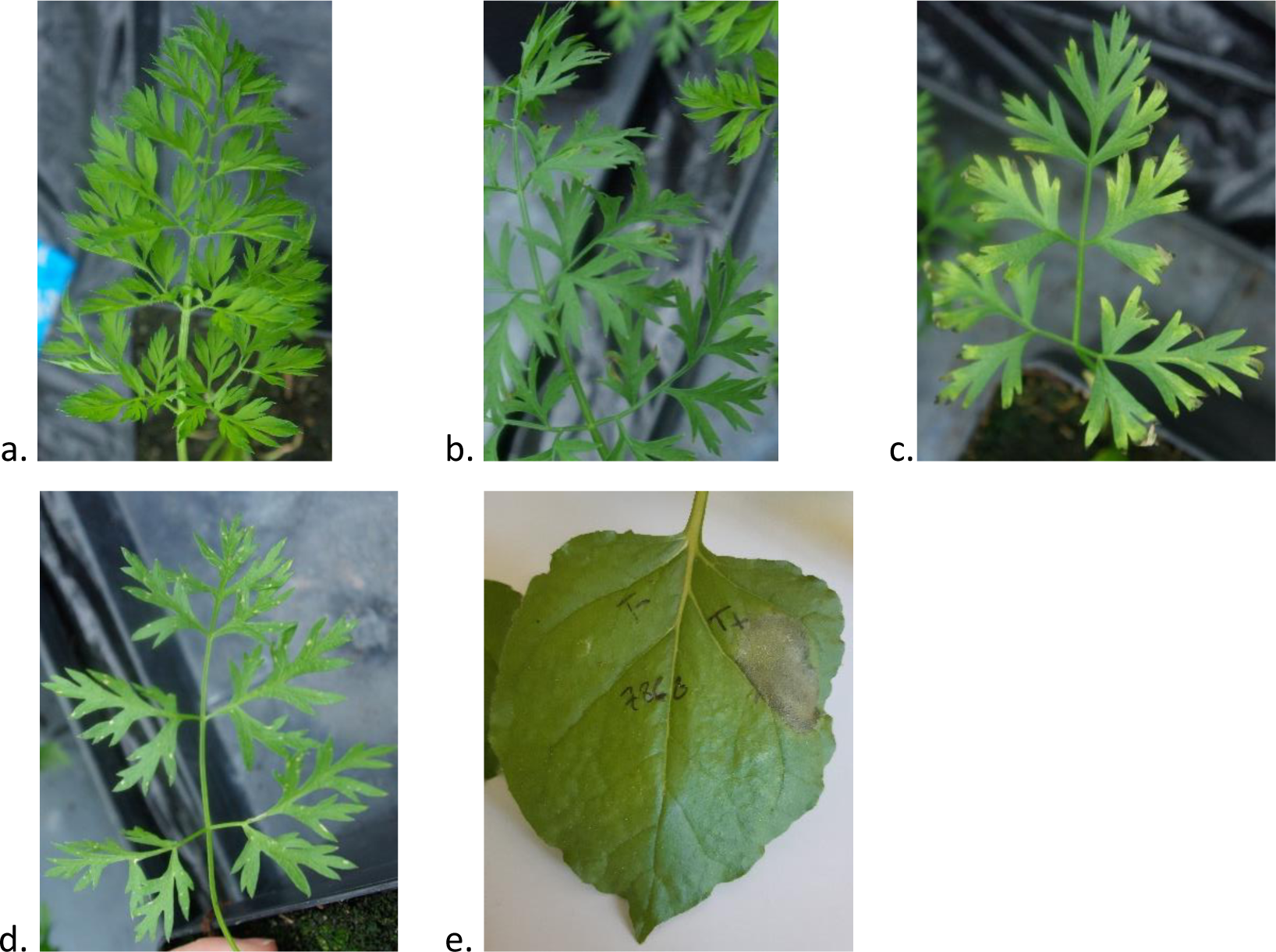
Pathogenicity test. a. b. c. d. on carrot leaves and e. on tobacco. a. absence of symptoms, b. and c. bacterial blight symptoms, d. and e. HR.

**Table 3:**
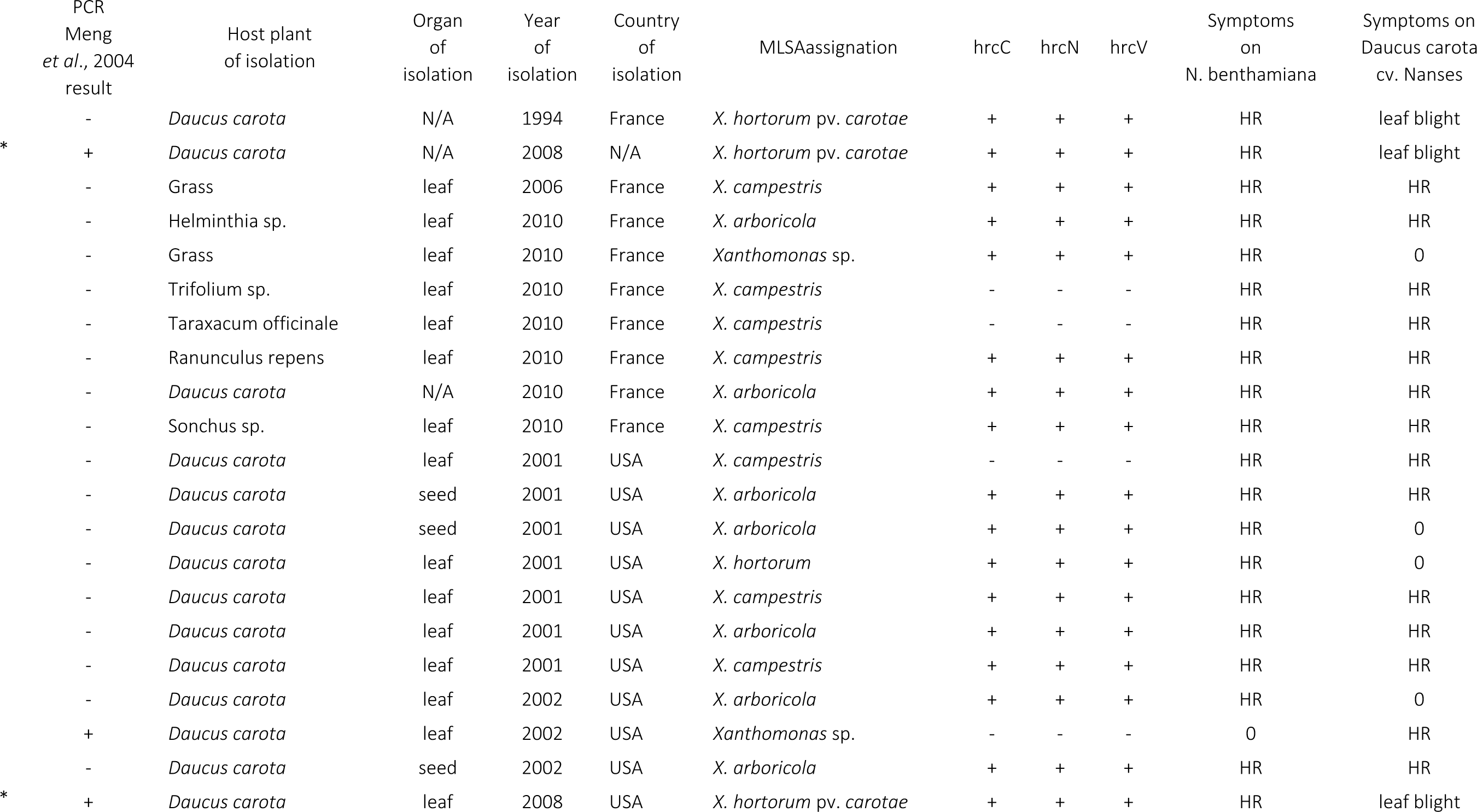
Search for the presence of hrc genes and pathogenicity on tobacco and carrot. *: positive control, HR: hypersensitive reaction, 0: absence of symptoms, leaf blight: typical symptoms of bacterial carrot leaf blight.

Pathogenic ability was also checked by the capacity of these 18 strains to induce an HR on *Nicotiana benthamiana*. Two- and three-days post inoculation, the 18 strains showed symptoms similar to those induced by the positive controls (CFBP 7768 and CFBP 7900), which were dry and non-evolving necrosis limited to the infiltrated area that are typical of an HR on tobacco (Table 3, Figure 2).

Finally, presence of genes coding core components (*hrcC, hrcN*, and *hrcV*) of an *hrp2*-type T3SS from the *Xanthomonas* group 2 were searched for by PCR (Merda et al., 2017). Typical signals were obtained for the three genes for 15 of these non-*X. hortorum* pv. *carotae* strains, but no signal was obtained for the most divergent strains that clustered close to *X. campestris* (CFBP 7806, CFBP 7807, CFBP 7840) (Table 3). It should, however, be indicated that the primers allow the amplification of the target genes in classical *X. campestris* strains (Merda et al., 2017). No signal was obtained for the *X. hortorum* pv. *carotae* strain CFBP 7868, which is coherent with its assignation within *Xanthomonas* group 1 that is known to present a non-canonical T3SS encoded by highly divergent genes (Wichmann et al., 2013).

From here, the strain CFBP 7706 was considered a typical *X. hortorum* pv. *carotae* strain together with all other 122 strains having led to a positive signal with Meng’s PCR test. Regarding strain CFBP 7868, while inducing a positive signal with Meng’s PCR test, the absence of pathogenicity on carrot and its distant phylogenetic position indicated that this strain does not belong to *X. hortorum* pv. *carotae*.

### 4.4. MLVA scheme development for getting insights in *X. hortorum* pv. *Carotae* genetic diversity

*In silico* analysis of the genome sequence of the strain M081 led to the identification of 14 TR loci (Table 2, Table S2). Eight of them (V1, V2, V3, V9, V10, V11, V12 and V14) demonstrated polymorphism on a subset of six strains, as indicated by bands of different sizes on agarose gel (Table 2). The six other TR loci (V4, V5, V6, V7, V8 and V13) were not amplified in all strains or did not show any polymorphism on agarose gel and were not further considered (Table S2). The eight selected VNTRs had TR unit size of 6 or 7 bp and a total fragment size ranging from 82 to 104 bp on the M081 genome. V14 is located on a coding region (SPOR domain-containing protein CDS), the four VNTRs V1, V3, V11, and V12 had only one of their primer and part of their offset fragment in coding regions (2OG-Fe(II) oxygenase CDS, hotdog fold thioesterase CDS, PH domain-containing protein CDS, *folB* CDS) and the three VNTRs V2, V9 and V10 were intergenic (Table 2).

All 122 *X. hortorum* pv. *carotae* strains were genotyped at all eight VNTR loci (Table 1). The MLVA-8 scheme was sufficient to reveal diversity in our dataset as it accurately discriminated the haplotypes in our strain collection, as the haplotype accumulation curve revealed that more than 95% of the haplotypes were detected with a set of seven markers (Figure 3).

**Figure 3:**
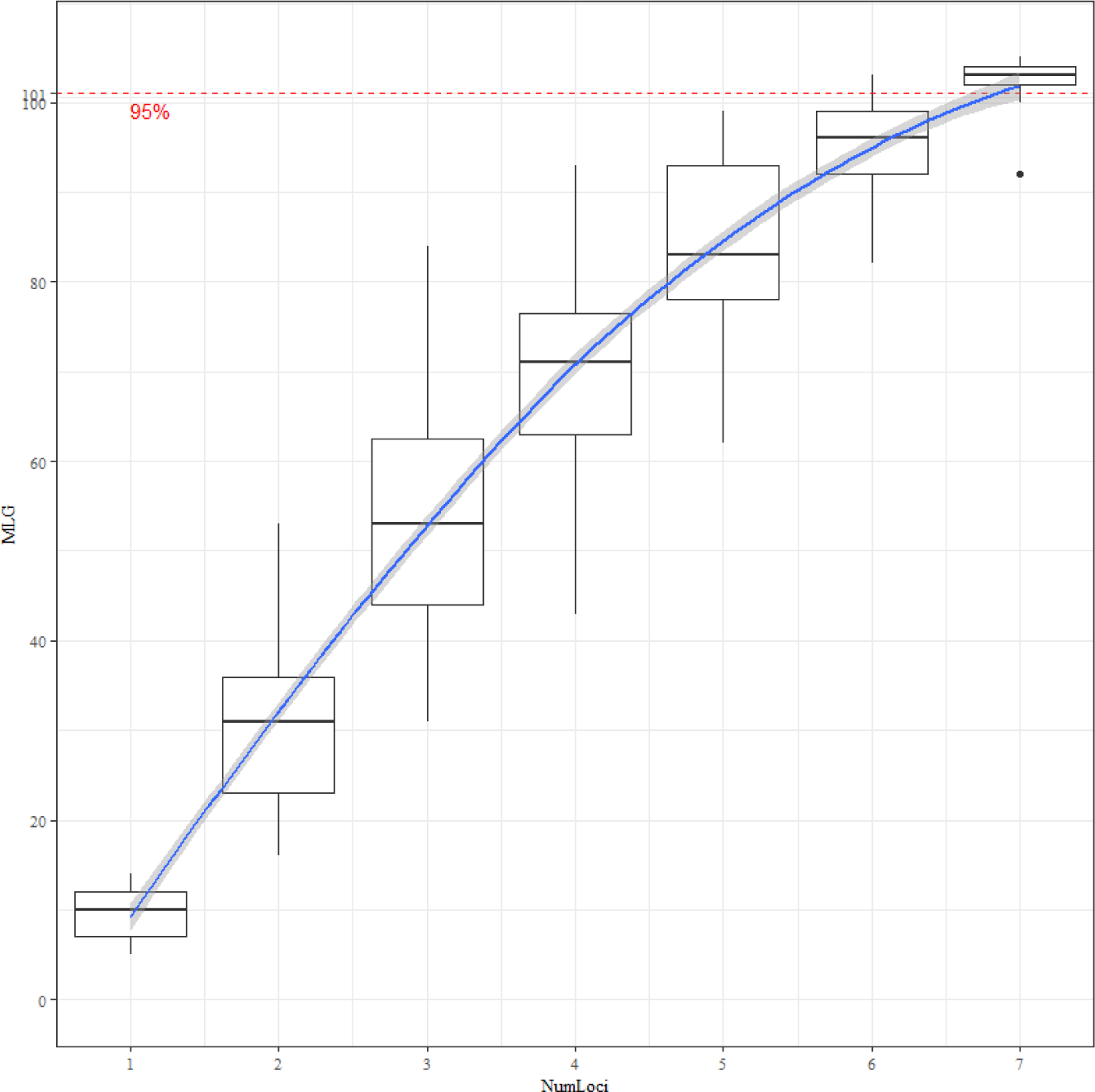
Genotype accumulation curve of *X. hortorum* pv. *carotae* over the 8 loci for the 122 strains of the study.

The MLVA 8-VNTR scheme allowed to observe a large diversity among the strains as they were typed in 106 haplotypes, with a discriminatory power of 0.9955, allowing further analyses. Diversity was also observed at each locus, with allele number per locus ranging from 5 to 14 and low Simpson’s diversity index ranging from 0.13 to 0.40 (Table 2).

DAPC analysis performed on the MLVA dataset allowed to group the strains into two clusters (Figure 4). Considering the position of the 26 strains previously characterized in MLSA, it appeared that these two groupings were identical (Figure 1). Among all alleles, 21 alleles were private to cluster A and 25 to cluster B (Table S3). At a whole, the 70 strains grouping in cluster A and 49 of the 52 strains grouping in cluster B had one to six private alleles, revealing strong differentiation between the two clusters (Table S3). A high allelic diversity was observed for both clusters, as the 70 strains of cluster A were typed in 61 haplotypes and the 52 strains of cluster B were typed in 45 haplotypes. Looking at the distribution of strains on Minimum Spanning Tree (MST), cluster A grouped the majority of the European strains (30/37) and approximatively one third of the North American ones (22/61). Cluster A grouped the strains into two clonal complexes (CC, i.e., networks grouping haplotypes differing from their closest neighbor at one or two VNTR loci) (Figure 5). The main CC grouped 58 of the 70 strains, typed in 49 haplotypes. The founder haplotype #99 of this main CC grouped seven strains isolated between 2005 and 2012, from France (4/7) and USA (1/7). The smaller CC grouped three strains isolated from different countries. The last nine strains were singletons that did not grouped in CC. Cluster B grouped 52 strains, the majority of which were isolated in the USA (39). Looking at the MST, strains from cluster B grouped in five CCs. The main CC was composed of 33 strains, typed in 28 haplotypes, from which the majority were isolated in the USA (27/33). The four small CC grouped two haplotypes, composed of one to two strains, that were isolated from similar area at the continental level, or were of unknown origin. It is interesting to note that the two strains isolated in Mauritius island formed a CC on their own and did not group with strains from other origins. Year of strain isolation was not a determinant of the grouping of strains in any cluster.

**Figure 4:**
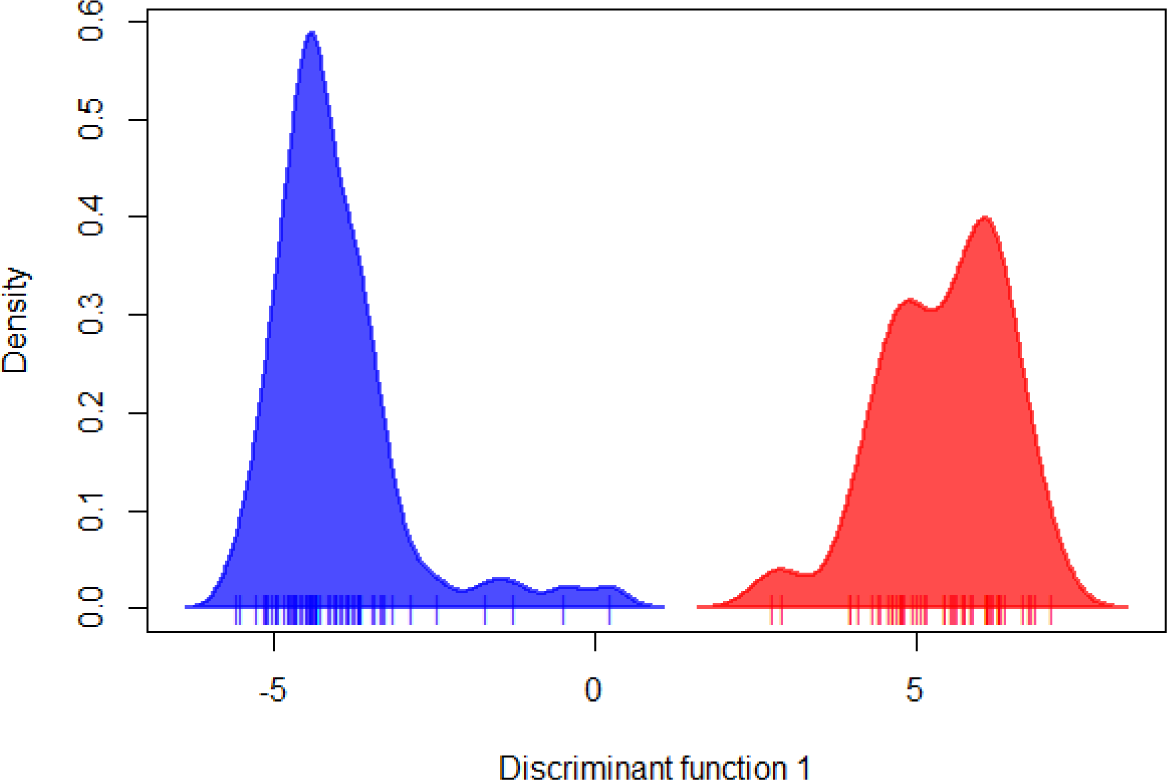
Discriminant Analysis of Principal Components clusterization of the 122 *X. hortorum* pv. *carotae* strains for k=2. Cluster blue grouped the strains from MLSA cluster A and cluster red the strains from MLSA cluster B.

**Figure 5:**
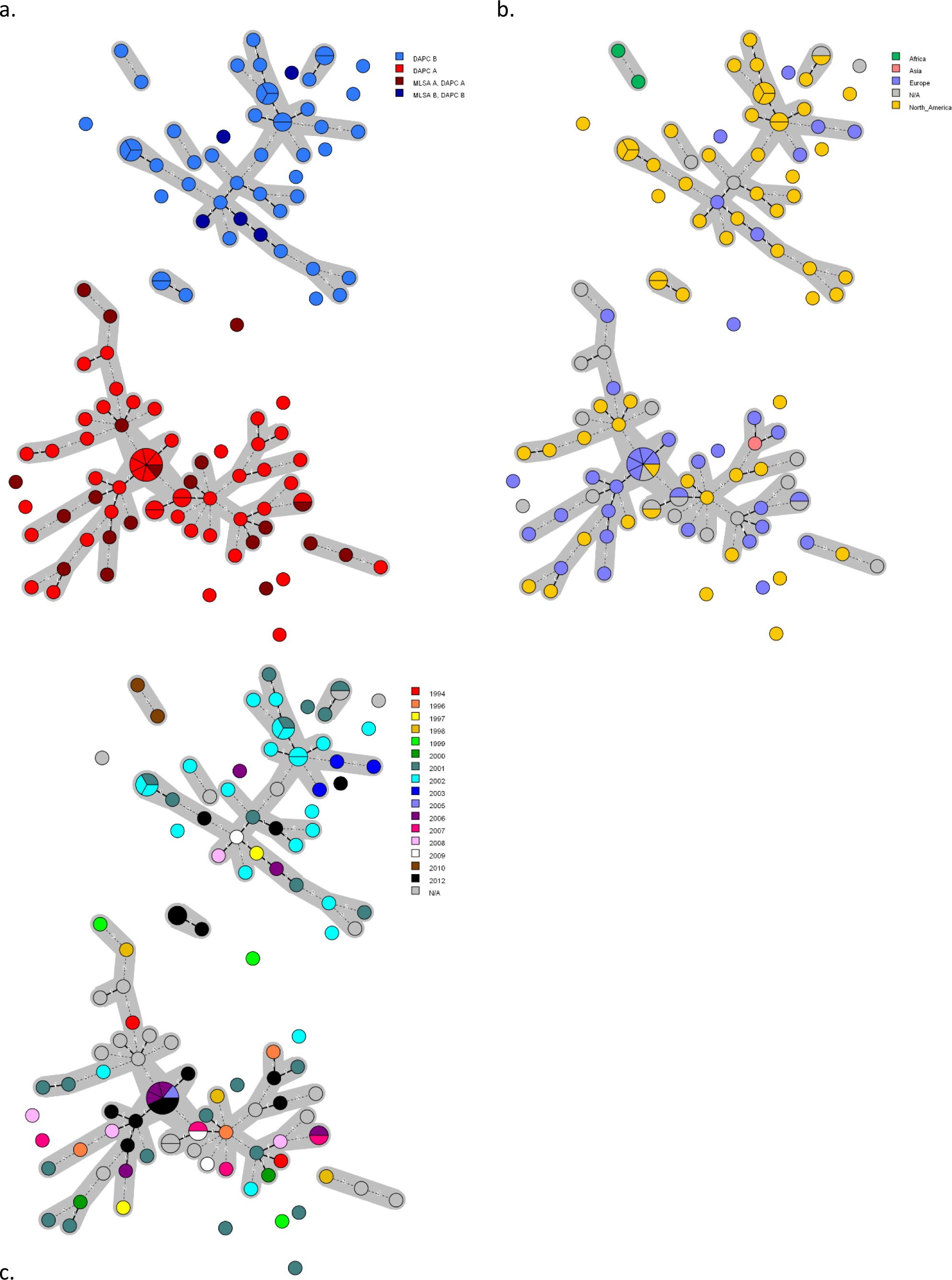
Minimum spanning tree of the 122 *X. hortorum* pv. *carotae* strains using VNTR-8 scheme. Dot diameter represents the number of strains per haplotype. Link number refer to the number of TR loci polymorphic and distinguishing two haplotypes. Grey area around haplotypes illustrate clonal complexes. Dot colors refer to a. MLSA and DAPC clusters; b. continent of sampling; c. year of sampling.

## 5. Discussion

*X. hortorum* pv. *carotae* is one of the three major pathogens infecting carrot crops and causing leaf blight (Groves and Skolko, 1944; Kimbrel et al., 2011). Found worldwide where carrot is cultivated, *X. hortorum* pv. *carotae* is a threat for this crop (du Toit et al., 2005). No efficient chemical treatment being available to cure *X. hortorum* pv. *carotae* -infected carrot plants (du Toit and Derie, 2008), control mostly relies on prophylactic measures. Being seed-transmitted, seed-health testing, seed thermotherapy, seed treatment and discarding infected seed-lots are currently the main solutions used to control this pathogen (du Toit et al., 2005). Knowledge on *X. hortorum* pv. *carotae* genetic structure is currently highly limited, but would be of great interest to inform genetic resistance studies of carrot to this pathogen and provide additional management option against this disease. Limited screening for resistance has so far been reported, but while no complete resistance was observed in the USDA carrot germplasm collection, substantial variation in severity of the disease was observed among the accessions that were tested indicating that breeding for resistance is an option to limit the impact of this disease (Christianson et al., 2015). Knowing the diversity of a pathogen is a prerequisite to test for resistance, because it allows to select the proper strains to be as representative as possible of this diversity and test lines against them all.

*X. hortorum* pv. *carotae* presents a low genetic diversity. Bacteria reproduce asexually by binary fission, but rely on various mechanisms of recombination to introduce some genetic diversity in their genome. Depending on the relative importance of mutation vs. recombination, bacterial species present a population structure ranging from strictly clonal to panmictic (Maynard-Smith et al., 1993). It has been previously shown for the species *Xanthomonas arboricola*, that bacterial plant pathogens have a clonal mode of evolution, while non-pathogenic strains form a recombinant network (Merda et al., 2016). Here, a MLSA based on seven housekeeping genes applied on 26 *X. hortorum* pv. *carotae* strains revealed a low genetic diversity within the pathovar *carotae*, as only 1.55% of sites were polymorphic (65 SNPs over the 4,189 bp analyzed). This pathovar is subdivided into two clusters, in which the 20 strains from cluster A differed from each other’s by only one SNP and all six strains from cluster B were identical. These strains were selected to maximize the diversity we gathered in our larger collection and were collected over 17 years, from seven countries and three continents. This low diversity revealed by MLSA indicates that clonal reproduction is dominant. The lack of resolution of MLSA to describe the genetic structure was previously observed for other pathogens of the Lysobacteraceae family (previously known as *Xanthomonadaceae*) (Almeida et al., 2010; Leduc et al., 2015; Ferreira et al., 2019; Dupas et al., 2023).

To overcome this problem, a MLVA scheme was developed and allowed us to type the entire collection of *X. hortorum* pv. *carotae* strains. The clusters A and B identified in MLSA were also observed in MLVA that, however, proved to be highly resolute as the 122 *X. hortorum* pv. *carotae* strains were typed in 106 haplotypes. Cluster A was composed of a majority of European strains and cluster B grouped the majority of American strains. This *X. hortorum* pv. *carotae* strain differentiation according to the geographical origin of the plant material can be linked to carrot types. Indeed, Europe and USA are known to mainly use different types of carrot genotypes. The Nantaise, Long Orange, Amsterdam or Paris types were typically bred and developed in Europe, while the Imperator or Danvers types are characteristic of the USA market (Stelmach et al., 2021). The seed production of Imperator type is mostly done in the USA, while seeds of Nantaise type are produced in Europe but also in the United States. This could suggest that strains infecting these two types of carrot develop mostly in allopatry and experience only limited gene flow, translating in the distinct groups revealed by MLSA and MLVA, except when two divergent carrot types are cropped close together, which can lead to mixing of strains, that are easily disseminated by air (du Toit et al., 2005; Scott and Dung, 2022).

A limited set of our *Xanthomonas*-like strains isolated from organs of symptomatic carrots or weeds (18/140) turned out to be non-*X. hortorum* pv. *carotae*, as mostly revealed by the use of the Meng’s PCR test (Meng et al., 2004) and confirmed by taxonomic assignation using MLSA. MLSA indicated that these non-pathogenic strains were genetically diverse and highly divergent from the pathogenic *X. hortorum* pv. *carotae* strains. Only one non-pathogenic strain belonged to the *X. hortorum* species, and all other scattered in the *Xanthomonas* phylogeny. Most non-*X. hortorum* pv. *carotae* strains isolated from carrot organs or weeds from *X. hortorum* pv. *carotae* infected carrot fields grouped with *X. arboricola* and *X. campestris*, which are already known to cluster non-pathogenic strains besides the classical pathovars described for a while (Cesbron et al., 2015; Merda et al., 2016; Lee et al., 2020). Most non-*X. hortorum* pv. *carotae* strains had all three *hrc* genes taken as indicators of a functional *hrp2* type-TTSS and these strains also induced an HR on tobacco. It is interesting to note that other strains that do not present a canonical *hrp2* type-T3SS do induce hypersensitive-like reactions on non-host plant, such as tobacco, a result indicating that these HRs are induced by other virulence factors (Meline et al., 2019).

We used the PCR-based identification test proposed by Meng and colleague to identify *X. hortorum* pv. *carotae* strains (Meng et al., 2004). This test proved highly useful, but presented one case of false positive and one case of a false negative results over the 44 tests we run (2.27% each). These results should be taken into account while using it in seed-health assays. By the way, it is important to highlight that the International Seed Federation protocol (ISHI-Veg, 2021) proposes to replace the gel-based Meng’s PCR test used in the ISTA protocol by a TaqMan test combining primers and probes from Barnhoorn 2014 (MVS*X. hortorum* pv. *carotae* 3 set) and from Temple *et al*. 2013 (*X. hortorum* pv. *carotae* -q2 set) (Temple et al., 2013; Barnhoorn, 2014).

### Proposition of strain CFBP 7900^NT^ as the neopathotype strain of *X. hortorum* pv. *carotae*

Since the holopathotype strain of *X. hortorum* pv. *carotae* is no longer available due to the storage of a contaminant (*Entoreobacter*) in international collections, we propose to replace it by the strain CFBP 7900, also known as M081, as the neopathotype strain of the pathovar *carotae* (Kimbrel et al., 2011). Strain CFBP 7900 was isolated in 2008 in a carrot plant in the USA, which is the same country than the *X. hortorum* pv. *carotae* holopathotype strain (Kendrick, 1934). The genome of this strain was previously sequenced and analyzed and is highly characteristic of that of a pathogenic *Xanthomonas* (Kimbrel et al., 2011). In our study, this strain was identified using the Meng’s PCR test (Meng et al., 2004) and the identification was confirmed by a taxonomic assignation using a MLSA based on seven housekeeping genes that included all other pathotype strains of pathovars of *X. hortorum*. This strain represents one of the two clusters forming the pathovar *carotae* (Figure 1). Moreover, the presence of 3 *hrc* genes was confirmed and indicate a functional TTSS, which is confirmed by the induction of an HR on tobacco. The strain was also found to be pathogenic on carrot and induced typical symptoms of leaf blight *e.i.* water-soaked lesions that evolved into necrotic spots, then dry and brittle, fitting initial description (Kendrick, 1934). Moreover, the strain CFBP 7900 was phenotypically characterized by standard biochemical tests and metabolic profiling on Biolog GEN III microplates (Morinière et al., 2020). All these data are consistent with the first description of the holopathotype made by Kendrick (Kendrick, 1934).

To date (04-03-2023) only three genomes of *X. hortorum* pv. *carotae* are publicly available in the National Center for Biotechnology Information (NCBI). Two of them are from the strain CFBP 7900 (= M081) and the third from a strain named MAFF 301101 isolated in Japan (BioSAmple: SAMN02470773, SAMEA6962650 and SAMN16925265) (Kimbrel et al., 2011; Dia et al., 2020). Only the first one deposited in 2013 and published by (Kimbrel et al., 2011) presents a high quality level and is circularized. Moreover, the strain M081 was deposited at the CIRM-CFBP, under the name CFBP 7900. It is publicly available and maintained by this international open collection.

The holopathotype strain of *X. hortorum* pv. *carotae* has been reported to be unsuitable as the pathotype strain of this pathovar, since at least 1991, leading to the use of other strains as reference when studying the pathovar *carotae* (Young et al., 1991). Thus, since 2011 and the genome sequencing of the strain CFBP 7900, it was used in a majority of studies designing molecular detection tests (Kimbrel et al., 2011; Temple et al., 2013; Dia et al., 2022), in studies using these molecular tests to confirm *X. hortorum* pv. *carotae* identification (Myung et al., 2014; Palomo et al., 2021), as a reference strain or reference genome to identify *X. hortorum* pv. *carotae* strains (Palomo et al., 2021), or in papers of other *Xanthomonas* species or pathovar as the reference for the pathovar *carotae* (Dhakal et al., 2019; Dia et al., 2021; Hébert et al., 2021; Morinière et al., 2021; Rosenthal et al., 2022).

## 6. Conclusion

This study revealed that *X. hortorum* pv. *carotae* is structured in two highly genetically homogeneous subgroups. These subgroups were revealed in MLSA and MLVA and may reflect allopatry of strains infecting different types of carrot genotypes, which is an important knowledge that could inform genetic breeding of carrot for resistance against the bacterial blight of carrot. This study provided genotyping data, which combined with data from the literature, leads to propose the strain CFBP 7900 (= M081) as the neopathotype strain of the pathovar *carotae*.

## 7. Author statements

### 7.1. Author contributions

Conceptualization: MAJ, ED, KD. Material and Methodology: MAJ, ED, KD. Supervision: MAJ. Writing-original draft: ED, MAJ. Writing-review & editing: ED, MAJ.

### 7.2. Conflicts of interest

*The authors declare that the research was conducted in the absence of any commercial or financial relationships that could be construed as a potential conflict of interest*.

## 7.3. Acknowledgements

We thanks Lindsey du Toit (Washington State University, USA),Olivier Pruvost (CIRAD), Isabelle Serandat (GEVES), Valérie Olivier (Anses), Perrine David (HM-Clause), Margaret Asma (Bejo), Benoit Meriaux (FNAMS) for strains donations. We also thank CIRM-CFBP (Beaucouzé, INRAE, France http://www6.inra.fr/cirm_eng/CFBP-Plant-Associated-Bacteria) for strain preservation and supply.

## Notes

### Competing Interest Statement

The authors have declared no competing interest.

