## Supplemental figures for "Analysis of the worldwide diversity of *Xanthomonas hortorum* pv. *carotae*, the agent of bacterial blight of carrot, reveals two distinct populations"

Figure S1: Phylogenetic relationships of the 26 *X. hortorum* pv. *carotae* and 18 non-*X. hortorum* pv. *carotae* strains from our collection evaluated by MLSA and compared to the pathotype strains of *X. hortorum* and the type strains of all species currently included in the genus *Xanthomonas*. Maximum likelihood tree was constructed on portion of *atpD* gene, for a total length of 738 bp, with bootstrap scores calculated on 100 replicates.

Figure S2: Phylogenetic relationships of the 26 *X. hortorum* pv. *carotae* and 18 non-*X. hortorum* pv. *carotae* strains from our collection evaluated by MLSA and compared to the pathotype strains of *X. hortorum* and the type strains of all species currently included in the genus *Xanthomonas*. Maximum likelihood tree was constructed on portion of *dnaK* gene, for a total length of 738 bp, with bootstrap scores calculated on 100 replicates.

Figure S3: Phylogenetic relationships of the 26 *X. hortorum* pv. *carotae* and 18 non-*X. hortorum* pv. *carotae* strains from our collection evaluated by MLSA and compared to the pathotype strains of *X. hortorum* and the type strains of all species currently included in the genus *Xanthomonas*. Maximum likelihood tree was constructed on portion of *efP* gene, for a total length of 327 bp, with bootstrap scores calculated on 100 replicates.

Figure S4: Phylogenetic relationships of the 26 *X. hortorum* pv. *carotae* and 18 non-*X. hortorum* pv. *carotae* strains from our collection evaluated by MLSA and compared to the pathotype strains of *X. hortorum* and the type strains of all species currently included in the genus *Xanthomonas*. Maximum likelihood tree was constructed on portion of *fyuA* gene, for a total length of 711 bp, with bootstrap scores calculated on 100 replicates.

Figure S5: Phylogenetic relationships of the 26 *X. hortorum* pv. *carotae* and 18 non-*X. hortorum* pv. *carotae* strains from our collection evaluated by MLSA and compared to the pathotype strains of *X. hortorum* and the type strains of all species currently included in the genus *Xanthomonas*. Maximum likelihood tree was constructed on portion of *glnA* gene, for a total length of 675 bp, with bootstrap scores calculated on 100 replicates.

Figure S6: Phylogenetic relationships of the 26 *X. hortorum* pv. *carotae* and 18 non-*X. hortorum* pv. *carotae* strains from our collection evaluated by MLSA and compared to the pathotype strains of *X. hortorum* and the type strains of all species currently included in the genus *Xanthomonas*. Maximum likelihood tree was constructed on portion of *gyrB* gene, for a total length of 627 bp, with bootstrap scores calculated on 100 replicates.

Figure S7: Phylogenetic relationships of the 26 *X. hortorum* pv. *carotae* and 18 non-*X. hortorum* pv. *carotae* strains from our collection evaluated by MLSA and compared to the pathotype strains of *X. hortorum* and the type strains of all species currently included in the genus *Xanthomonas*. Maximum likelihood tree was constructed on portion of *rpoD* gene, for a total length of 389 bp, with bootstrap scores calculated on 100 replicates.

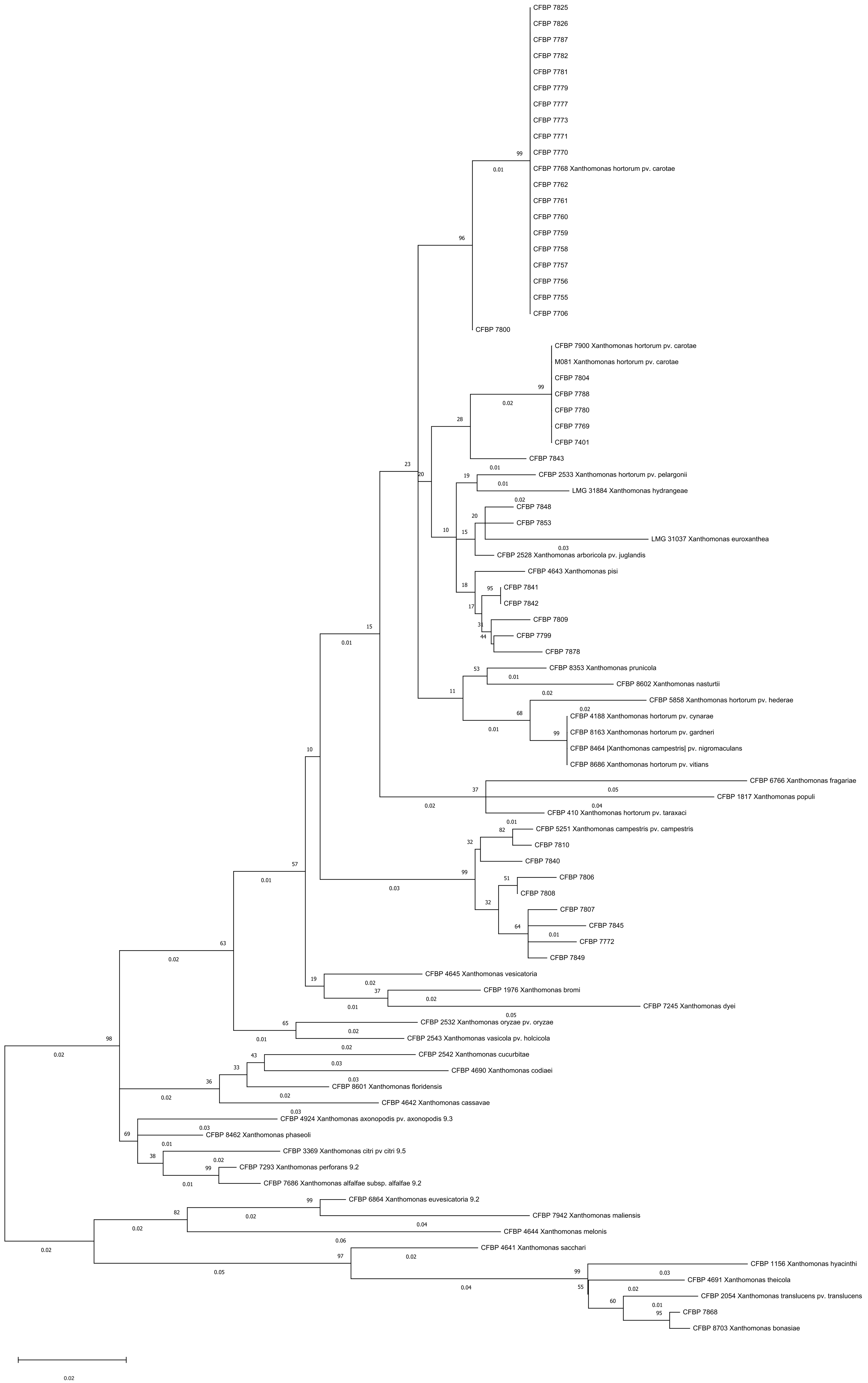
CFBP 7825
CFBP 7826
CFBP 7787
CFBP 7782
CFBP 7781
CFBP 7779
CFBP 7777
CFBP 7773
CFBP 7771
CFBP 7770
CFBP 7768 Xanthomonas hortorum pv. carotae
CFBP 7762
CFBP 7761
CFBP 7760
CFBP 7759
CFBP 7758
CFBP 7757
CFBP 7756
CFBP 7755
CFBP 7706
CFBP 7800
CFBP 7900 Xanthomonas hortorum pv. carotae
M081 Xanthomonas hortorum pv. carotae
CFBP 7804
CFBP 7788
CFBP 7780
CFBP 7769
CFBP 7401
CFBP 7843
CFBP 2533 Xanthomonas hortorum pv. pelargonii
LMG 31884 Xanthomonas hydrangeae
CFBP 7848
CFBP 7853
LMG 31037 Xanthomonas euroxanthea
CFBP 2528 Xanthomonas arboricola pv. juglandis
CFBP 4643 Xanthomonas pisi
CFBP 7841
CFBP 7842
CFBP 7809
CFBP 7799
CFBP 7878
CFBP 8353 Xanthomonas prunicola
CFBP 8602 Xanthomonas nasturtii
CFBP 5858 Xanthomonas hortorum pv. hederae
CFBP 4188 Xanthomonas hortorum pv. cynarae
CFBP 8163 Xanthomonas hortorum pv. gardneri
CFBP 8464 |Xanthomonas campestris| pv. nigromaculans
CFBP 8686 Xanthomonas hortorum pv. vitians
CFBP 6766 Xanthomonas fragariae
CFBP 1817 Xanthomonas populi
CFBP 410 Xanthomonas hortorum pv. taraxaci
CFBP 5251 Xanthomonas campestris pv. campestris
CFBP 7810
CFBP 7840
CFBP 7806
CFBP 7808
CFBP 7807
CFBP 7845
CFBP 7772
CFBP 7849
CFBP 4645 Xanthomonas vesicatoria
CFBP 1976 Xanthomonas bromi
CFBP 7245 Xanthomonas dyei
CFBP 2532 Xanthomonas oryzae pv. oryzae
CFBP 2543 Xanthomonas vasicola pv. holcicola
CFBP 2542 Xanthomonas cucurbitae
CFBP 4690 Xanthomonas codiaei
CFBP 8601 Xanthomonas floridensis
CFBP 4642 Xanthomonas cassavae
CFBP 4924 Xanthomonas axonopodis pv. axonopodis 9.3
CFBP 8462 Xanthomonas phaseoli
CFBP 3369 Xanthomonas citri pv citri 9.5
CFBP 7293 Xanthomonas perforans 9.2
CFBP 7686 Xanthomonas alfalfae subsp. alfalfae 9.2
CFBP 6864 Xanthomonas euvesicatoria 9.2
CFBP 7942 Xanthomonas maliensis
CFBP 4644 Xanthomonas melonis
CFBP 4641 Xanthomonas sacchari
CFBP 1156 Xanthomonas hyacinthi
CFBP 4691 Xanthomonas theicola
CFBP 2054 Xanthomonas translucens pv. translucens
CFBP 7868
CFBP 8703 Xanthomonas bonasiae
0.02

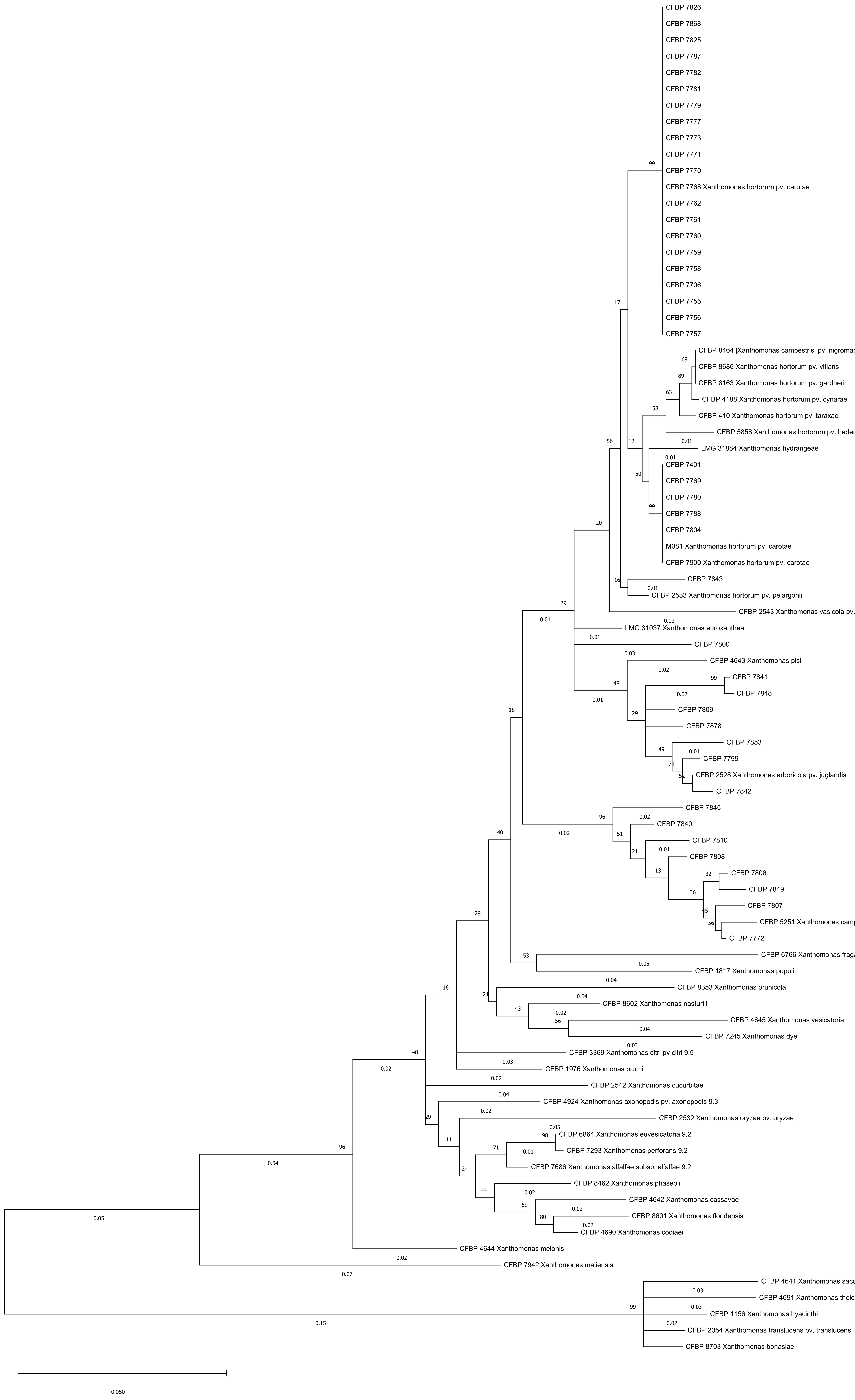
CFBP 7826
CFBP 7868
CFBP 7825
CFBP 7787
CFBP 7782
CFBP 7781
CFBP 7779
CFBP 7777
CFBP 7773
CFBP 7771
CFBP 7770
CFBP 7768 Xanthomonas hortorum pv. carotae
CFBP 7762
CFBP 7761
CFBP 7760
CFBP 7759
CFBP 7758
CFBP 7706
CFBP 7755
CFBP 7756
CFBP 7757
CFBP 8464 |Xanthomonas campestris| pv. nigromaculans
CFBP 8686 Xanthomonas hortorum pv. vitians
CFBP 8163 Xanthomonas hortorum pv. gardneri
CFBP 4188 Xanthomonas hortorum pv. cynarae
CFBP 410 Xanthomonas hortorum pv. taraxaci
CFBP 5858 Xanthomonas hortorum pv. hederae
LMG 31884 Xanthomonas hydrangeae
CFBP 7401
CFBP 7769
CFBP 7780
CFBP 7788
CFBP 7804
M081 Xanthomonas hortorum pv. carotae
CFBP 7900 Xanthomonas hortorum pv. carotae
CFBP 7843
CFBP 2533 Xanthomonas hortorum pv. pelargonii
CFBP 2543 Xanthomonas vasicola pv. holcicola
LMG 31037 Xanthomonas euroxanthea
CFBP 7800
CFBP 4643 Xanthomonas pisi
CFBP 7841
CFBP 7848
CFBP 7809
CFBP 7878
CFBP 7853
CFBP 7799
CFBP 2528 Xanthomonas arboricola pv. juglandis
CFBP 7842
CFBP 7845
CFBP 7840
CFBP 7810
CFBP 7808
CFBP 7806
CFBP 7849
CFBP 7807
CFBP 5251 Xanthomonas campestris pv. campestris
CFBP 7772
CFBP 6766 Xanthomonas fragariae
CFBP 1817 Xanthomonas populi
CFBP 8353 Xanthomonas prunicola
CFBP 8602 Xanthomonas nasturtii
CFBP 4645 Xanthomonas vesicatoria
CFBP 7245 Xanthomonas dyei
CFBP 3369 Xanthomonas citri pv citri 9.5
CFBP 1976 Xanthomonas bromi
CFBP 2542 Xanthomonas cucurbitae
CFBP 4924 Xanthomonas axonopodis pv. axonopodis 9.3
CFBP 2532 Xanthomonas oryzae pv. oryzae
CFBP 6864 Xanthomonas euvesicatoria 9.2
CFBP 7293 Xanthomonas perforans 9.2
CFBP 7686 Xanthomonas alfalfae subsp. alfalfae 9.2
CFBP 8462 Xanthomonas phaseoli
CFBP 4642 Xanthomonas cassavae
CFBP 8601 Xanthomonas floridensis
CFBP 4690 Xanthomonas codiaei
CFBP 4644 Xanthomonas melonis
CFBP 7942 Xanthomonas maliensis
CFBP 4641 Xanthomonas sacchari
CFBP 4691 Xanthomonas theicola
CFBP 1156 Xanthomonas hyacinthi
CFBP 2054 Xanthomonas translucens pv. translucens
CFBP 8703 Xanthomonas bonasiae
0.050

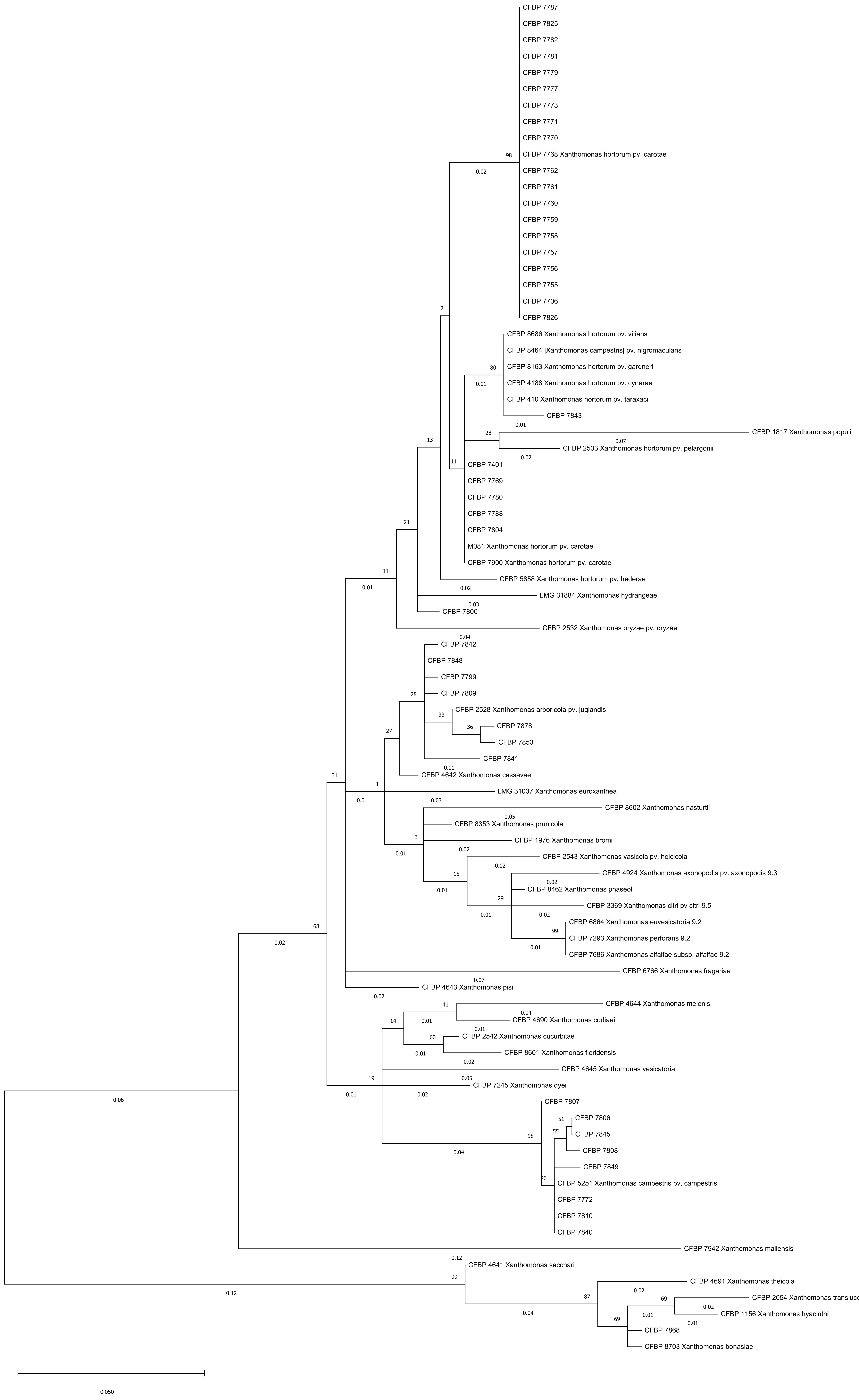

CFBP 7787
CFBP 7825
CFBP 7782
CFBP 7781
CFBP 7779
CFBP 7777
CFBP 7773
CFBP 7771
CFBP 7770
CFBP 7768 Xanthomonas hortorum pv. carotae
CFBP 7762
CFBP 7761
CFBP 7760
CFBP 7759
CFBP 7758
CFBP 7757
CFBP 7756
CFBP 7755
CFBP 7706
CFBP 7826
CFBP 8686 Xanthomonas hortorum pv. vitians
CFBP 8464 |Xanthomonas campestris| pv. nigromaculans
CFBP 8163 Xanthomonas hortorum pv. gardneri
CFBP 4188 Xanthomonas hortorum pv. cynarae
CFBP 410 Xanthomonas hortorum pv. taraxaci
CFBP 7843
CFBP 1817 Xanthomonas populi
CFBP 2533 Xanthomonas hortorum pv. pelargonii
CFBP 7401
CFBP 7769
CFBP 7780
CFBP 7788
CFBP 7804
M081 Xanthomonas hortorum pv. carotae
CFBP 7900 Xanthomonas hortorum pv. carotae
CFBP 5858 Xanthomonas hortorum pv. hederae
LMG 31884 Xanthomonas hydrangeae
CFBP 7800
CFBP 2532 Xanthomonas oryzae pv. oryzae
CFBP 7842
CFBP 7848
CFBP 7799
CFBP 7809
CFBP 2528 Xanthomonas arboricola pv. juglandis
CFBP 7878
CFBP 7853
CFBP 7841
CFBP 4642 Xanthomonas cassavae
LMG 31037 Xanthomonas euroxanthea
CFBP 8602 Xanthomonas nasturtii
CFBP 8353 Xanthomonas prunicola
CFBP 1976 Xanthomonas bromi
CFBP 2543 Xanthomonas vasicola pv. holcicola
CFBP 4924 Xanthomonas axonopodis pv. axonopodis 9.3
CFBP 8462 Xanthomonas phaseoli
CFBP 3369 Xanthomonas citri pv citri 9.5
CFBP 6864 Xanthomonas euvesicatoria 9.2
CFBP 7293 Xanthomonas perforans 9.2
CFBP 7686 Xanthomonas alfalfae subsp. alfalfae 9.2
CFBP 6766 Xanthomonas fragariae
CFBP 4643 Xanthomonas pisi
CFBP 4644 Xanthomonas melonis
CFBP 4690 Xanthomonas codiaei
CFBP 2542 Xanthomonas cucurbitae
CFBP 8601 Xanthomonas floridensis
CFBP 4645 Xanthomonas vesicatoria
CFBP 7245 Xanthomonas dyei
CFBP 7807
CFBP 7806
CFBP 7845
CFBP 7808
CFBP 7849
CFBP 5251 Xanthomonas campestris pv. campestris
CFBP 7772
CFBP 7810
CFBP 7840
CFBP 7942 Xanthomonas maliensis
CFBP 4641 Xanthomonas sacchari
CFBP 4691 Xanthomonas theicola
CFBP 2054 Xanthomonas translucens pv. translucens
CFBP 1156 Xanthomonas hyacinthi
CFBP 7868
CFBP 8703 Xanthomonas bonasiae
0.050

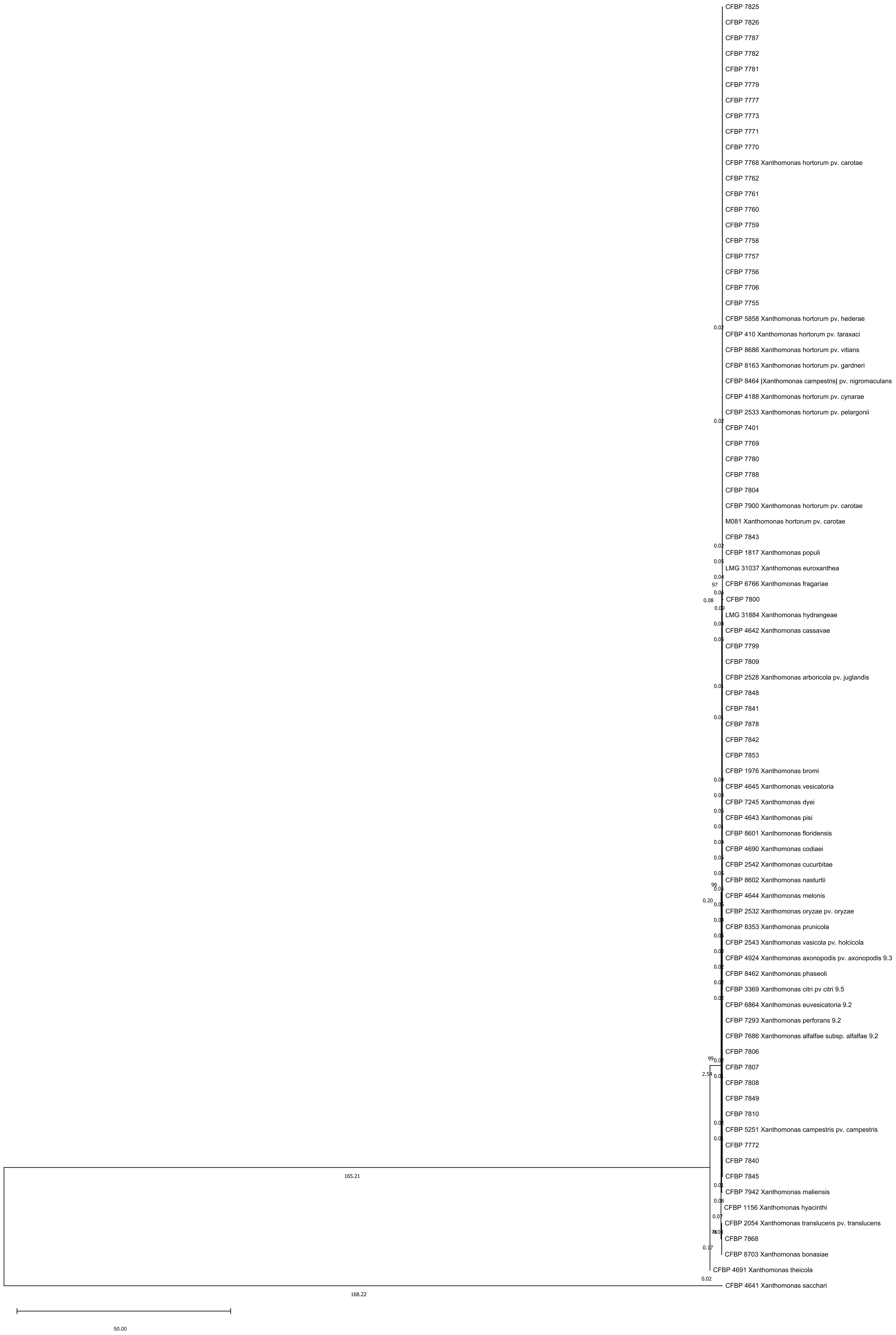
CFBP 7825
CFBP 7826
CFBP 7787
CFBP 7782
CFBP 7781
CFBP 7779
CFBP 7777
CFBP 7773
CFBP 7771
CFBP 7770
CFBP 7768 Xanthomonas hortorum pv. carotae
CFBP 7762
CFBP 7761
CFBP 7760
CFBP 7759
CFBP 7758
CFBP 7757
CFBP 7756
CFBP 7706
CFBP 7755
CFBP 5858 Xanthomonas hortorum pv. hederae
0.02
CFBP 410 Xanthomonas hortorum pv. taraxaci
CFBP 8686 Xanthomonas hortorum pv. vitians
CFBP 8163 Xanthomonas hortorum pv. gardneri
CFBP 8464 |Xanthomonas campestris| pv. nigromaculans
CFBP 4188 Xanthomonas hortorum pv. cynarae
CFBP 2533 Xanthomonas hortorum pv. pelargonii
0.02
CFBP 7401
CFBP 7769
CFBP 7780
CFBP 7788
CFBP 7804
CFBP 7900 Xanthomonas hortorum pv. carotae
M081 Xanthomonas hortorum pv. carotae
CFBP 7843
0.02
CFBP 1817 Xanthomonas populi
LMG 31037 Xanthomonas euroxanthea
97
0.08
CFBP 6766 Xanthomonas fragariae
CFBP 7800
LMG 31884 Xanthomonas hydrangeae
CFBP 4642 Xanthomonas cassavae
CFBP 7799
CFBP 7809
CFBP 2528 Xanthomonas arboricola pv. juglandis
CFBP 7848
CFBP 7841
CFBP 7878
CFBP 7842
CFBP 7853
CFBP 1976 Xanthomonas bromi
CFBP 4645 Xanthomonas vesicatoria
CFBP 7245 Xanthomonas dyei
CFBP 4643 Xanthomonas pisi
CFBP 8601 Xanthomonas floridensis
CFBP 4690 Xanthomonas codiaei
CFBP 2542 Xanthomonas cucurbitae
CFBP 8602 Xanthomonas nasturtii
99
0.20
CFBP 4644 Xanthomonas melonis
CFBP 2532 Xanthomonas oryzae pv. oryzae
CFBP 8353 Xanthomonas prunicola
CFBP 2543 Xanthomonas vasicola pv. holcicola
CFBP 4924 Xanthomonas axonopodis pv. axonopodis 9.3
CFBP 8462 Xanthomonas phaseoli
CFBP 3369 Xanthomonas citri pv citri 9.5
CFBP 6864 Xanthomonas euvesicatoria 9.2
CFBP 7293 Xanthomonas perforans 9.2
CFBP 7686 Xanthomonas alfalfae subsp. alfalfae 9.2
CFBP 7806
99
CFBP 7807
CFBP 7808
CFBP 7849
CFBP 7810
CFBP 5251 Xanthomonas campestris pv. campestris
CFBP 7772
CFBP 7840
165.21
CFBP 7845
CFBP 7942 Xanthomonas maliensis
0.08
CFBP 1156 Xanthomonas hyacinthi
0.07
CFBP 2054 Xanthomonas translucens pv. translucens
CFBP 7868
0.17
CFBP 8703 Xanthomonas bonasiae
CFBP 4691 Xanthomonas theicola
0.02
CFBP 4641 Xanthomonas sacchari
168.22
50.00

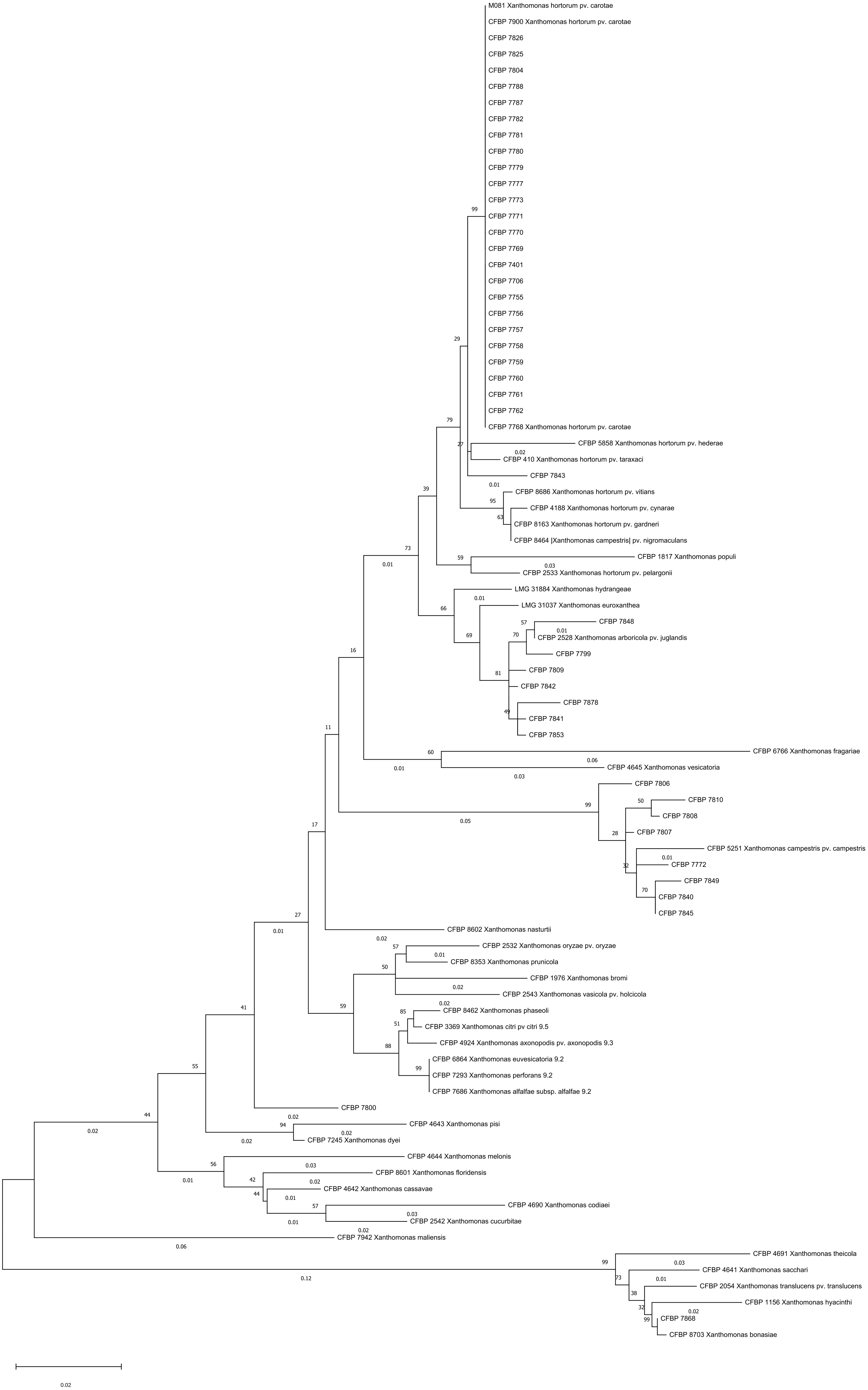
M081 Xanthomonas hortorum pv. carotae
CFBP 7900 Xanthomonas hortorum pv. carotae
CFBP 7826
CFBP 7825
CFBP 7804
CFBP 7788
CFBP 7787
CFBP 7782
CFBP 7781
CFBP 7780
CFBP 7779
CFBP 7777
CFBP 7773
CFBP 7771
CFBP 7770
CFBP 7769
CFBP 7401
CFBP 7706
CFBP 7755
CFBP 7756
CFBP 7757
CFBP 7758
CFBP 7759
CFBP 7760
CFBP 7761
CFBP 7762
CFBP 7768 Xanthomonas hortorum pv. carotae
CFBP 5858 Xanthomonas hortorum pv. hederae
CFBP 410 Xanthomonas hortorum pv. taraxaci
CFBP 7843
CFBP 8686 Xanthomonas hortorum pv. vitians
CFBP 4188 Xanthomonas hortorum pv. cynarae
CFBP 8163 Xanthomonas hortorum pv. gardneri
CFBP 8464 |Xanthomonas campestris| pv. nigromaculans
CFBP 1817 Xanthomonas populi
CFBP 2533 Xanthomonas hortorum pv. pelargonii
LMG 31884 Xanthomonas hydrangeae
LMG 31037 Xanthomonas euroxanthea
CFBP 7848
CFBP 2528 Xanthomonas arboricola pv. juglandis
CFBP 7799
CFBP 7809
CFBP 7842
CFBP 7878
CFBP 7841
CFBP 7853
CFBP 6766 Xanthomonas fragariae
CFBP 4645 Xanthomonas vesicatoria
CFBP 7806
CFBP 7810
CFBP 7808
CFBP 7807
CFBP 5251 Xanthomonas campestris pv. campestris
CFBP 7772
CFBP 7849
CFBP 7840
CFBP 7845
CFBP 8602 Xanthomonas nasturtii
CFBP 2532 Xanthomonas oryzae pv. oryzae
CFBP 8353 Xanthomonas prunicola
CFBP 1976 Xanthomonas bromi
CFBP 2543 Xanthomonas vasicola pv. holcicola
CFBP 8462 Xanthomonas phaseoli
CFBP 3369 Xanthomonas citri pv citri 9.5
CFBP 4924 Xanthomonas axonopodis pv. axonopodis 9.3
CFBP 6864 Xanthomonas euvesicatoria 9.2
CFBP 7293 Xanthomonas perforans 9.2
CFBP 7686 Xanthomonas alfalfae subsp. alfalfae 9.2
CFBP 7800
CFBP 4643 Xanthomonas pisi
CFBP 7245 Xanthomonas dyei
CFBP 4644 Xanthomonas melonis
CFBP 8601 Xanthomonas floridensis
CFBP 4642 Xanthomonas cassavae
CFBP 4690 Xanthomonas codiaei
CFBP 2542 Xanthomonas cucurbitae
CFBP 7942 Xanthomonas maliensis
CFBP 4691 Xanthomonas theicola
CFBP 4641 Xanthomonas sacchari
CFBP 2054 Xanthomonas translucens pv. translucens
CFBP 1156 Xanthomonas hyacinthi
CFBP 7868
CFBP 8703 Xanthomonas bonasiae
0.02

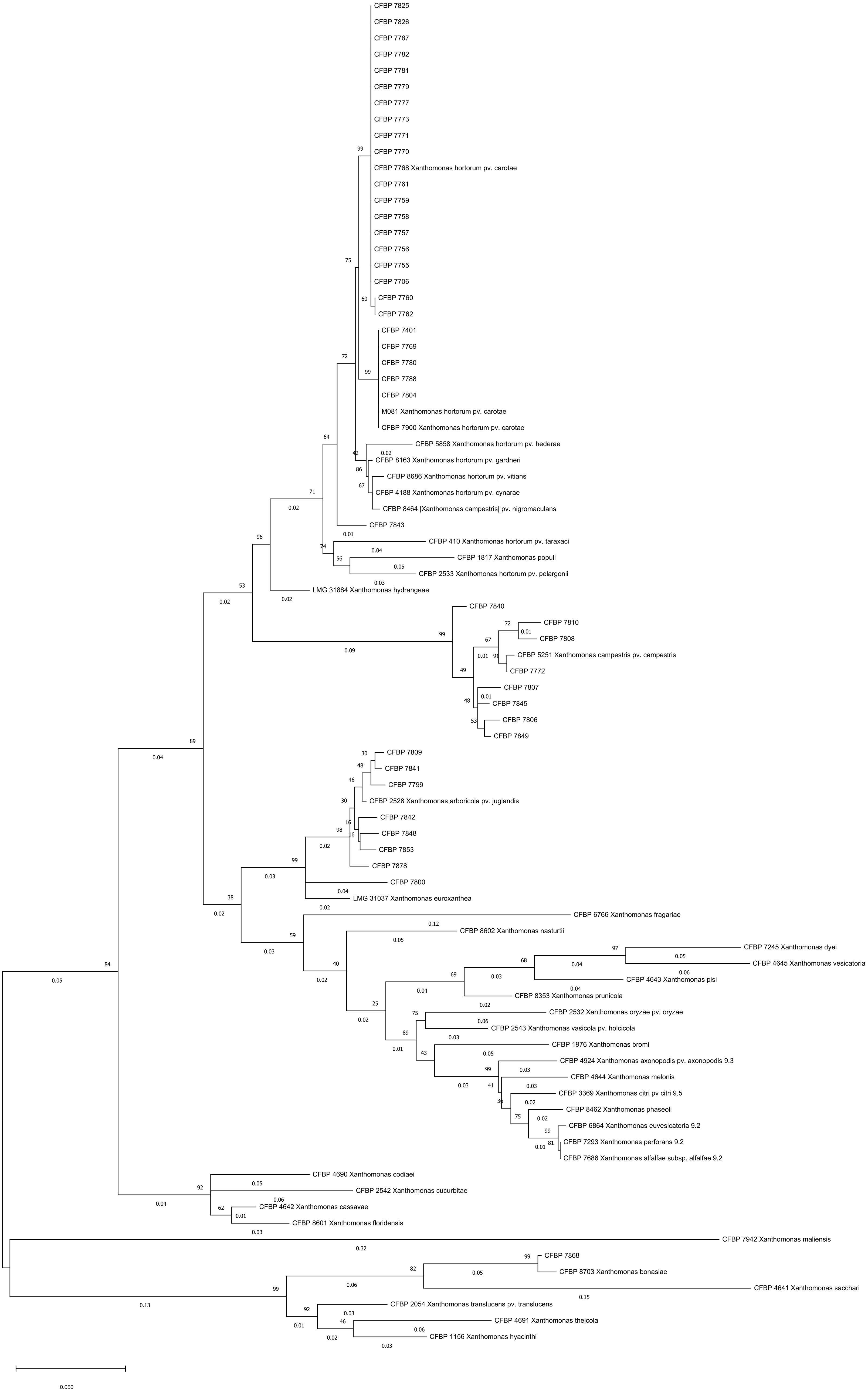
CFBP 7825
CFBP 7826
CFBP 7787
CFBP 7782
CFBP 7781
CFBP 7779
CFBP 7777
CFBP 7773
CFBP 7771
CFBP 7770
CFBP 7768 Xanthomonas hortorum pv. carotae
CFBP 7761
CFBP 7759
CFBP 7758
CFBP 7757
CFBP 7756
CFBP 7755
CFBP 7706
CFBP 7760
CFBP 7762
CFBP 7401
CFBP 7769
CFBP 7780
CFBP 7788
CFBP 7804
M081 Xanthomonas hortorum pv. carotae
CFBP 7900 Xanthomonas hortorum pv. carotae
CFBP 5858 Xanthomonas hortorum pv. hederae
CFBP 8163 Xanthomonas hortorum pv. gardneri
CFBP 8686 Xanthomonas hortorum pv. vitians
CFBP 4188 Xanthomonas hortorum pv. cynarae
CFBP 8464 |Xanthomonas campestris| pv. nigromaculans
CFBP 7843
CFBP 410 Xanthomonas hortorum pv. taraxaci
CFBP 1817 Xanthomonas populi
CFBP 2533 Xanthomonas hortorum pv. pelargonii
LMG 31884 Xanthomonas hydrangeae
CFBP 7840
CFBP 7810
CFBP 7808
CFBP 5251 Xanthomonas campestris pv. campestris
CFBP 7772
CFBP 7807
CFBP 7845
CFBP 7806
CFBP 7849
CFBP 7809
CFBP 7841
CFBP 7799
CFBP 2528 Xanthomonas arboricola pv. juglandis
CFBP 7842
CFBP 7848
CFBP 7853
CFBP 7878
CFBP 7800
LMG 31037 Xanthomonas euroxanthea
CFBP 6766 Xanthomonas fragariae
CFBP 8602 Xanthomonas nasturtii
CFBP 7245 Xanthomonas dyei
CFBP 4645 Xanthomonas vesicatoria
CFBP 4643 Xanthomonas pisi
CFBP 8353 Xanthomonas prunicola
CFBP 2532 Xanthomonas oryzae pv. oryzae
CFBP 2543 Xanthomonas vasicola pv. holcicola
CFBP 1976 Xanthomonas bromi
CFBP 4924 Xanthomonas axonopodis pv. axonopodis 9.3
CFBP 4644 Xanthomonas melonis
CFBP 3369 Xanthomonas citri pv citri 9.5
CFBP 8462 Xanthomonas phaseoli
CFBP 6864 Xanthomonas euvesicatoria 9.2
CFBP 7293 Xanthomonas perforans 9.2
CFBP 7686 Xanthomonas alfalfae subsp. alfalfae 9.2
CFBP 4690 Xanthomonas codiaei
CFBP 2542 Xanthomonas cucurbitae
CFBP 4642 Xanthomonas cassavae
CFBP 8601 Xanthomonas floridensis
CFBP 7942 Xanthomonas maliensis
CFBP 7868
CFBP 8703 Xanthomonas bonasiae
CFBP 4641 Xanthomonas sacchari
CFBP 2054 Xanthomonas translucens pv. translucens
CFBP 4691 Xanthomonas theicola
CFBP 1156 Xanthomonas hyacinthi
0.050

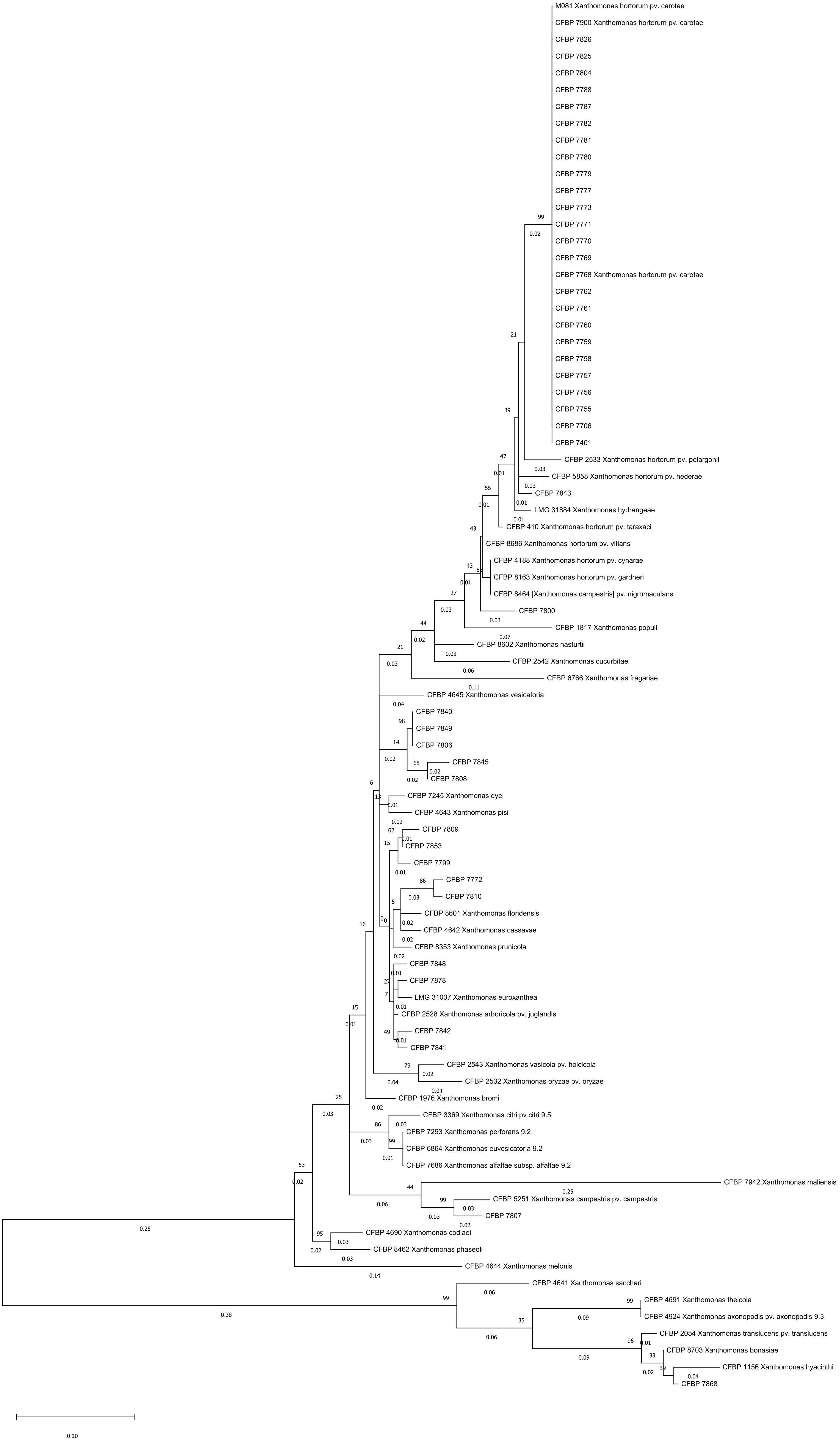
M081 Xanthomonas hortorum pv. carotae
CFBP 7900 Xanthomonas hortorum pv. carotae
CFBP 7826
CFBP 7825
CFBP 7804
CFBP 7788
CFBP 7787
CFBP 7782
CFBP 7781
CFBP 7780
CFBP 7779
CFBP 7777
CFBP 7773
CFBP 7771
CFBP 7770
CFBP 7769
CFBP 7768 Xanthomonas hortorum pv. carotae
CFBP 7762
CFBP 7761
CFBP 7760
CFBP 7759
CFBP 7758
CFBP 7757
CFBP 7756
CFBP 7755
CFBP 7706
CFBP 7401
CFBP 2533 Xanthomonas hortorum pv. pelargonii
CFBP 5858 Xanthomonas hortorum pv. hederae
CFBP 7843
LMG 31884 Xanthomonas hydrangeae
CFBP 410 Xanthomonas hortorum pv. taraxaci
CFBP 8686 Xanthomonas hortorum pv. vitians
CFBP 4188 Xanthomonas hortorum pv. cynarae
CFBP 8163 Xanthomonas hortorum pv. gardneri
CFBP 8464 |Xanthomonas campestris| pv. nigromaculans
CFBP 7800
CFBP 1817 Xanthomonas populi
CFBP 8602 Xanthomonas nasturtii
CFBP 2542 Xanthomonas cucurbitae
CFBP 6766 Xanthomonas fragariae
CFBP 4645 Xanthomonas vesicatoria
CFBP 7840
CFBP 7849
CFBP 7806
CFBP 7845
CFBP 7808
CFBP 7245 Xanthomonas dyei
CFBP 4643 Xanthomonas pisi
CFBP 7809
CFBP 7853
CFBP 7799
CFBP 7772
CFBP 7810
CFBP 8601 Xanthomonas floridensis
CFBP 4642 Xanthomonas cassavae
CFBP 8353 Xanthomonas prunicola
CFBP 7848
CFBP 7878
LMG 31037 Xanthomonas euroxanthea
CFBP 2528 Xanthomonas arboricola pv. juglandis
CFBP 7842
CFBP 7841
CFBP 2543 Xanthomonas vasicola pv. holcicola
CFBP 2532 Xanthomonas oryzae pv. oryzae
CFBP 1976 Xanthomonas bromi
CFBP 3369 Xanthomonas citri pv citri 9.5
CFBP 7293 Xanthomonas perforans 9.2
CFBP 6864 Xanthomonas euvesicatoria 9.2
CFBP 7686 Xanthomonas alfalfae subsp. alfalfae 9.2
CFBP 7942 Xanthomonas maliensis
CFBP 5251 Xanthomonas campestris pv. campestris
CFBP 7807
CFBP 4690 Xanthomonas codiaei
CFBP 8462 Xanthomonas phaseoli
CFBP 4644 Xanthomonas melonis
CFBP 4641 Xanthomonas sacchari
CFBP 4691 Xanthomonas theicola
CFBP 4924 Xanthomonas axonopodis pv. axonopodis 9.3
CFBP 2054 Xanthomonas translucens pv. translucens
CFBP 8703 Xanthomonas bonasiae
CFBP 1156 Xanthomonas hyacinthi
CFBP 7868
0.10
